# A decrease in fatty acid synthesis rescues cells with limited peptidoglycan synthesis capacity

**DOI:** 10.1101/2022.12.03.519008

**Authors:** Jessica R. Willdigg, Yesha Patel, John D. Helmann

**Author notes:** Address correspondence to John D. Helmann. Competing Interest Statement: The authors declare no conflict of interest.

## Abstract

Proper synthesis and maintenance of a multilayered cell envelope is critical for bacterial fitness. However, whether mechanisms exist to coordinate synthesis of the membrane and peptidoglycan layers is unclear. In *Bacillus subtilis*, synthesis of peptidoglycan (PG) during cell elongation is mediated by an elongasome complex acting in concert with class A PBPs (aPBPs). We previously described mutant strains limited in their capacity for PG synthesis due to a loss of aPBPs and an inability to compensate by upregulation of elongasome function. Growth of these PG-limited cells can be restored by suppressor mutations predicted to decrease membrane synthesis. One suppressor mutation leads to an altered function repressor, FapR*\**, that functions as a super-repressor and leads to decreased transcription of fatty acid synthesis (FAS) genes. Consistent with fatty acid limitation mitigating cell wall synthesis defects, inhibition of FAS by cerulenin also restored growth of PG-limited cells. Moreover, cerulenin can counteract the inhibitory effect of β-lactams in some strains. These results imply that limiting PG synthesis results in impaired growth, in part, due to an imbalance of PG and cell membrane synthesis and that *B. subtilis* lacks a robust physiological mechanism to reduce membrane synthesis when PG synthesis is impaired.

**Importance:** Understanding how a bacterium coordinates cell envelope synthesis is essential to fully appreciate how bacteria grow, divide, and resist cell envelope stresses such as β-lactam antibiotics. Balanced synthesis of the peptidoglycan cell wall and the cell membrane is critical for cells to maintain shape, turgor pressure and resist external cell envelope threats. Using *Bacillus subtilis*, we show that cells deficient in peptidoglycan synthesis can be rescued by compensatory mutations that decrease the synthesis of fatty acids. Further, we show that inhibiting fatty acid synthesis with cerulenin is sufficient to restore growth of cells deficient in peptidoglycan synthesis. Understanding the coordination of cell wall and membrane synthesis may provide insights relevant to antimicrobial treatment.

## Introduction

The balanced synthesis of the peptidoglycan (PG) cell wall, the cell membrane, and any additional envelope layers is important for the cell to maintain shape and resist turgor pressure and external cell envelope threats (1, 2). Many of our most powerful antibiotics target PG synthesis, and others disrupt membrane synthesis or integrity. Understanding the consequences of antibiotic inhibition, and the processes that ultimately lead to growth cessation or cell death, will facilitate efforts to counter the emerging threat of antibiotic resistance.

The PG cell wall is an important structure for nearly all bacteria, allowing cells to resist internal turgor pressure to prevent lysis as well as conferring shape and rigidity to the cell. Seemingly contrary to its role in maintaining rigidity, the cell wall is a highly dynamic structure (1, 2). The PG sacculus is constantly remodeled to accommodate the insertion of newly synthesized glycan strands during cell growth and division (3). Lipid II, a lipid-linked disaccharide pentapeptide PG precursor, is flipped across the cell membrane, and then the glycopeptide subunits are incorporated into the sacculus by penicillin-binding proteins (PBPs) (4). The bifunctional class A PBPs (aPBPs) contain both transglycosylase (TG) activity to incorporate new disaccharide units into the growing PG chains and transpeptidase (TP) activity to cross-link the adjacent strands. The class B PBPs (bPBPs) have TP activity, but lack TG function. An initially puzzling study revealed that *Bacillus subtilis* cells lacking all four aPBPs (Δ4 aPBP) still synthesize PG (5), which implied the presence of an alternative TG enzyme for assembly of the glycan strands. Subsequent work revealed that RodA provides TG activity to a multiprotein complex known as the elongasome, where it functions together with bPBPs that provide TP activity (6–8).

Elongation of the rod-shaped *B. subtilis* cell results from the balanced action of the elongasome and aPBPs (Figure 1A). The elongasome complex includes RodA (TG), bPBPs (TP), RodZ, MreC, and MreD, and is associated with a cytosolic filament comprising MreB and its paralogs (MreBH, Mbl). This elongasome complex moves circumferentially along the short axis of the cell and is associated with endopeptidases, including the MreBH-associated LytE protein (Figure 1A). Elongasome function is also thought to be supported by, or to help sustain, regions of increased membrane fluidity known as RIFs (9). The aPBPs move independently and diffusely during PG assembly (10–13). Although these PG synthesis machines do not move in a coordinated manner, both aPBPs and the elongasome are necessary to maintain normal cell length and width (Figure 1A) (14, 15).

**Figure 1.**
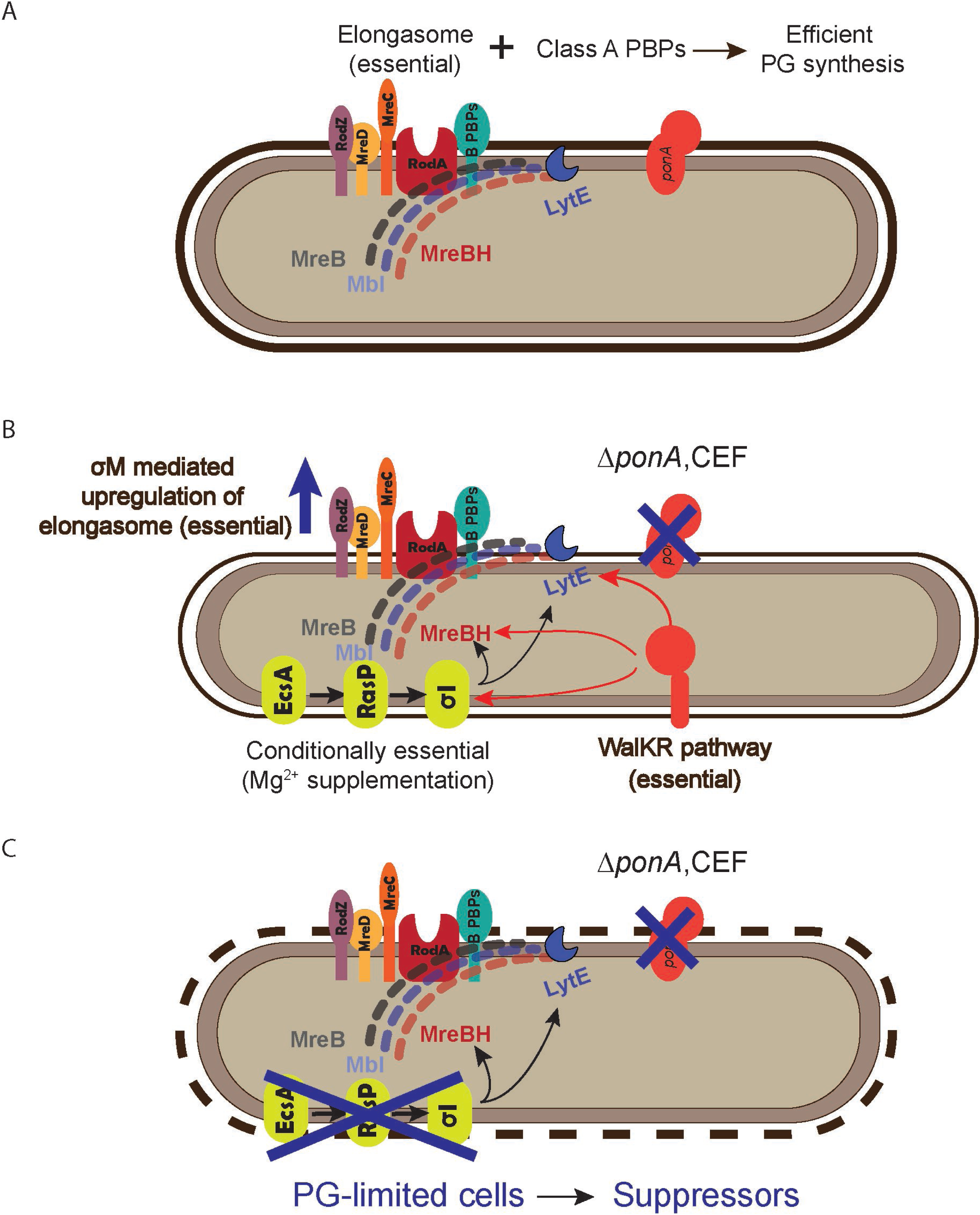
A model of PG-limitation in *B. subtilis* cells. (A) Efficient PG synthesis requires the coordination of the class A PBPs (aPBPs) and the essential elongasome protein complex. (B). Elimination of *ponA*-mediated PG synthesis (via either deletion of chemical inhibition by CEF) requires compensatory changes in elongasome gene expression for optimal PG synthesis. This is accomplished by upregulation of the essential TGase RodA (by σ^M^) and by upregulation of the conditionally essential *ecsA, rasP, sigI* pathway that increases expression of *mreBH* and *lytE.* Increased elongasome activity leads to thinner and elongated cells (15). (C). Cells that are deficient in *ponA* activity and unable to appropriately upregulate σ^I^ (*ecsA ponA, rasP ponA* and *sigI ponA* double mutants), are PG-limited and are conditionally lethal in the absence of Mg^2+^ supplementation. Mutations in the essential *walKR* pathway may arise that increase *mreBH* and *lytE* expression (19). Here, we focus on a spontaneous suppressor mutation (*fapR*\*) that restores growth on LB in the absence of Mg^2+^ by reduction in FAS.

Despite recent advances, there is still much to learn about PG synthesis and assembly. *B. subtilis* has served as the premier model system for the study of envelope synthesis in Gram-positive bacteria. We have previously characterized several cell envelope stress response (CESR) pathways that facilitate adaptation to antibiotics that inhibit envelope synthesis or function (16). For example, the extracytoplasmic function (ECF) sigma factor σ^M^ is necessary for intrinsic resistance to PG synthesis inhibitors (17, 18). In circumstances where the class A PBPs (aPBPs) are absent due to mutation or have been inhibited by antibiotics, cells depend on the elongasome for PG synthesis (Figure 1B). Several components of the elongasome, including RodA, are transcriptionally activated by σ^M^ in response to cell envelope stress (6).

Previously, we used TnSeq to identify genes that become essential in cells deficient for aPBP activity (19). This led to the elucidation of a regulatory pathway that leads to the σ^I^-dependent upregulation of the LytE endopeptidase and MreBH scaffolding protein (19). This pathway is essential for growth of cells deficient for aPBPs on LB medium (Figure 1B). However, growth on LB is rescued if the medium is supplemented with 20 mM magnesium (Mg^2+^) (19), as seen also for other mutant strains with defects in PG synthesis (20). The mechanism of this suppression is unknown but may be related to a reduction in autolysin activity (20).

The σ^I^ pathway is initiated by the EcsA membrane protein, which is required for the function of the RasP intramembrane protease (21). RasP, in turn, cleaves multiple proteins including the anti-sigma factors for σ^V^ (22, 23), σ^W^ (21, 24), and σ^I^ (25), which each coordinate a cell envelope stress response (26, 27). In cells lacking all four aPBPs (Δ4 aPBP), or even just the dominant enzyme PBP1 (*ponA*), the loss of *ecsA*, *rasP*, or *sigI* prevents growth on LB medium (19). These cells fail to grow since they are unable to activate σ^I^ to upregulate elongasome activity to compensate for the loss of aPBP function. We refer to the phenotypically similar *ecsA ponA*, *rasP ponA*, and *sigI ponA* double mutants as PG-limited strains (Figure 1C).

Here, we took advantage of the conditional lethality of PG-limited strains (*ecsA ponA* and *rasP ponA)* to select for growth in the absence of high Mg^2+^. This selection revealed a cohort of suppressor mutations in genes important for fatty acid synthesis (FAS). One of these mutants, *fapR\**, encodes a super-repressor that downregulates the expression of FAS enzymes. Further, we show that inhibiting FAS with cerulenin (CER) is sufficient to restore growth of PG-limited mutant strains. Characterization of the *fapR\** suppressor mutation highlights the benefits of balancing membrane and PG synthesis during cell envelope biogenesis. Moreover, these genetic findings suggest that, at least in this model system, cells lack a robust physiological feedback mechanism to reduce membrane synthesis in PG-limited cells, and that imbalanced synthesis contributes to fitness defects.

## Results

### Spontaneous mutations in FAS genes compensate for lethal cell wall defects in *B. subtilis*

We selected for mutations that restore growth on LB of strains deficient in peptidoglycan (PG) synthesis due to defects in both the major aPBP (*ponA*) and a pathway that would normally up-regulate elongasome function (*ecsA, rasP,* or *sigI*) (Figure 1C). For reference, we include a summary list of genes relevant to this work in Table S1. In previous work, we described suppressor mutations in *walK*, encoding an essential two-component system kinase. The WalK kinase and associated WalR transcription factor control the expression of autolysins and factors affecting autolytic activity. Thus, a gain-of-function mutation in WalK can increase expression of the MreBH and LytE proteins and substitute for the defective σ^I^ pathway (19). Here, we report the recovery of suppressor mutations in genes affecting FAS. Suppressor mutations were identified in the genes for all four subunits of the essential enzyme acetyl-CoA carboxylase (ACC), which catalyzes the carboxylation of acetyl-CoA to malonyl-CoA, and constitutes the first committed step of FAS (28) (Table 1). We hypothesize that these mutations encode reduced function alleles of ACC that restrict the capacity for membrane biosynthesis. We also recovered both missense and deletion mutations in *fapR*, encoding the fatty acid and phospholipid synthesis repressor FapR (29). FapR responds to accumulation of malonyl-CoA as a ligand to de-repress the synthesis of fatty acids and membrane phospholipids when this precursor is abundant (30).

**Table 1.** Spontaneous suppressor mutations recovered in PG-limited cells. Mutants of the ecsA ponA and rasP ponA parent strains were selected that were able to grow on LB medium in the absence of Mg^2+^ supplementation. Each suppressor strain contained a single point mutation identified via whole genome re-sequencing. The genome position was determined via comparison to the B. subtills reference genome NC_000964.3.

| Isolate | Strain Background | Gene | Genome Position | Nucleotide Change | Mutation |
| --- | --- | --- | --- | --- | --- |
| HB23854 | *ecsA ponA* | *fapR* | 1662276 | A --> G | Δ24 bp deletion in CDS/ I104V |
| HB23855 | *ecsA ponA* | *fapR* | 1662301 | C --> T | A112V |
| HB23856 | *ecsA ponA* | *accA* | 2988324 | C --> A | G129W |
| HB23857 | *ecsA ponA* | *accA* | 2988009 | A --> G | T234R |
| HB23860 | *ecsA ponA* | *accB* | 2532222 | T --> C | SNP between promoter and RBS |
| HB23885 | *rasP ponA* | *fapR* | 1662190 | T --> A | 175N |
| HB23874 | *rasP ponA* | *accA* | 2988055 | G --> T | N218K |
| HB23875 | *rasP ponA* | *accC* | 2531202 | T --> C | R169G |
| HB23883 | *rasP ponA* | *accC* | 2531393 | G --> A | P105L |
| HB23877 | *rasP ponA* | *accD* | 2989081 | G --> A | T162M |
| HB23879 | *rasP ponA* | *accD* | 2989654 | T --> deletion | deletion in promoter |

### *fapR*\* is a dominant mutation that rescues growth but not cell morphology

We first focused on understanding the mechanism by which an in-frame *fapR* deletion mutation, here defined as *fapR*\* (Figure 2A), suppressed the growth defect of PG-limited parent strains. A strain lacking the major aPBP (*ponA*) in combination with either *ecsA*, *rasP*, or *sigI* is unable to grow in LB medium. However, *fapR*\* rescues growth of all three double mutant strains (Figure 2B). In contrast with *fapR*\*, deletion of *fapR* did not rescue growth of *sigI ponA* (Figure 2C). This suggests that *fapR\** is an altered function mutation. These findings are consistent with our previous data (19), which place *ecsA*, *rasP*, and *sigI* in a regulatory pathway that upregulates both *mreBH* and *lytE* in support of elongasome function (Figure 1B). In this pathway, the RasP protease cleaves the RsgI protein, a membrane-embedded anti-sigma factor containing and intrinsicially disordered region postulated to sense cell wall integrity (25).

**Figure 2.**
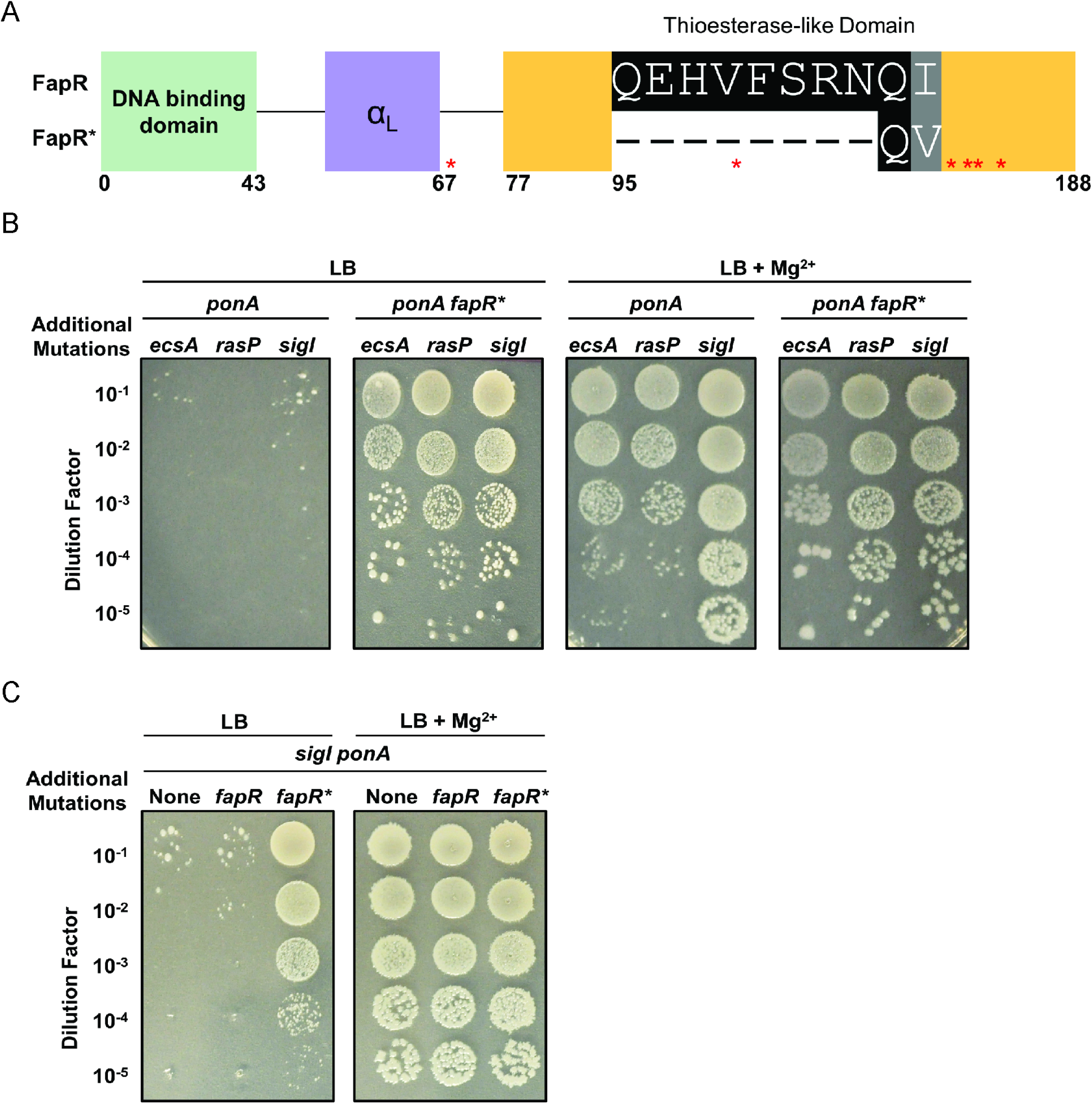
The *fapR*\* suppressor mutation rescues cells with PG synthesis defects. (A) The *fapR\** 24-nucleotide deletion leads to an 8 amino acid deletion in the thioesterase-like, malonyl-CoA binding domain of the FapR repressor. Asterisks indicate FapR residues important for coordinating malonyl-CoA. (B) *fapR\** rescues viability of *ecsA ponA*, *rasP ponA*, and *sigI ponA* cells in LB medium in the absence of Mg^2+^. (C) Deletion of *fapR* does not restore viability of a *sigI ponA* mutant grown in LB medium in the absence of Mg^2+^.

Next, we sought to determine how *fapR\** affects cell growth and morphology in both the wild-type (WT) background and in suppressed strains. Compared to WT, *fapR\** strains had a significantly slower average doubling time in LB medium (WT, 24 ± 0.03 min, *fapR*,* 39 ± 0.11 min) (Figure S1A). Moreover, ectopic overexpression of *fapR\** (but not *fapR*) resulted in slower growth in a merodiploid strain, indicating that *fapR*\* is a dominant mutation (Figure S1B). Although slower growing, the *fapR\** mutant was unaltered in shape as judged by microscopic analysis of cell length and width (Figure S2). In contrast, the PG-limited *rasP ponA* cells transferred to LB medium lacking added Mg^2+^ displayed a variety of morphological abnormalities including examples of filamentation, bending, and coiling (Figure S3A). These abnormalities were partially suppressed in the Mg^2+^-amended medium, but some lysed ghost cells were still visible (Figure S3B). Although introduction of the *fapR*\* allele rescued growth in LB, it did not restore a normal cell morphology to the PG-limited strain. In the *rasP ponA fapR*\* strain we observed numerous examples of filamented cells, many with enhanced coiling (Figure S3C). Thus, abnormal morphology seems to be a poor proxy for understanding the growth limitations that occur in these PG-limited cells.

### The *fapR\** mutation downregulates the FapR regulon

The *fapR*\* allele contains a 24-nucleotide deletion in the thioesterase-like domain overlapping with residue Phe99, known to be necessary for binding of the malonyl-CoA inducer (Figure 2A) (30). Thus, we hypothesized that FapR* might have reduced sensitivity to malonyl-CoA, which would lead to tighter repression of FAS genes. To test this hypothesis, we measured sensitivity to cerulenin (CER). CER is a 12-carbon mycotoxin that inhibits FAS by forming a covalent linkage with β-ketoacyl-ACP synthases, including FabF, to block fatty acid elongation (31–33). Upon inhibition by CER, malonyl-CoA accumulates, triggering de-repression of genes in the FapR regulon (34). Since *fapR\** is postulated to reduce the sensitivity of FapR to malonyl-CoA, we asked whether *fapR*\* mutants were more sensitive to CER. As predicted, *fapR*\* mutant strains had greatly increased sensitivity to CER at concentrations that did not inhibit either the WT or *fapR* null mutant strains (Figure 3A). This effect was specific to CER and unlikely to reflect defects in general membrane permeability since no change was noted in sensitivity to other common hydrophobic antibiotics (Figure S4).

**Figure 3.**
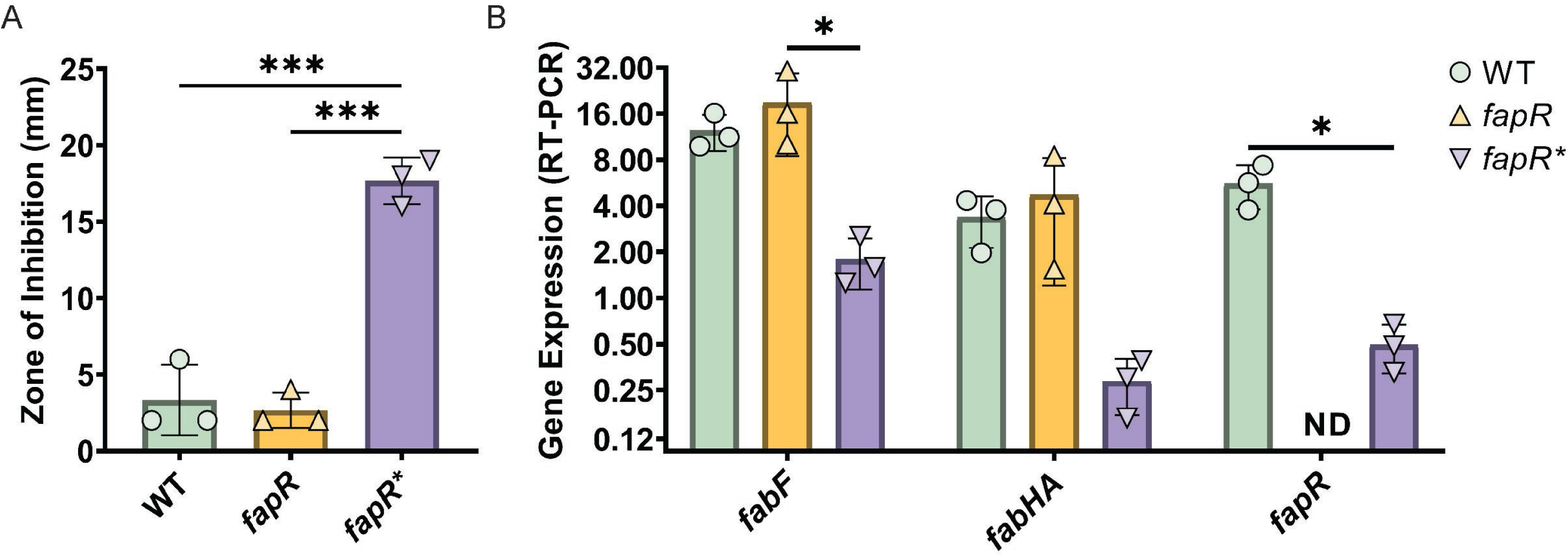
*fapR\** decreases the expression of FAS genes. (A) *fapR\** exhibits an increased sensitivity to cerulenin (CER) compared to WT and *fapR* null cells. 10 µg CER was applied to the disc (6mm), and disc diameter was subtracted from zone of inhibition values. A one-way ANOVA with Tukey test for multiple comparisons was performed. *p*-value cutoff (***) < 0.0002. (B) Gene expression from representative genes in the FapR regulon (*fabF, fapR and fabHA)*. The legend indicates strain genotypes. ND indicates the expected lack of detectable *fapR* mRNA in a *fapR* null strain. Gene expression values (2^-Δct^) were normalized to *gyrA*. A one-way ANOVA with a Tukey test for multiple comparisons was performed. *p*-value cutoff (*) < 0.05.

We next used qRT-PCR to directly probe the impact of FapR* on the expression levels of selected genes in the FapR regulon. In the *fapR* null strain, *fapR* was not detected (as expected) and expression of *fabF* and *fabHA* genes was changed little, although the data are consistent with a modest increase (Figure 3B). This suggests that FapR may be largely inactive as a repressor in these mid-logarithmic phase cells. In contrast, we noted a 5-10-fold decrease in the expression levels of *fabF, fabHA,* and *fapR* in the *fapR*\* strain (Figure 3B). Thus, the *fapR*\* allele results in a higher level of repression of genes within the FapR regulon.

### FapR* is a super-repressor with reduced response to malonyl-CoA

We hypothesized that FapR*\** functions as a dominant super-repressor due to a diminished ability to sense malonyl-CoA as an inducer. FapR recognizes a 17-base pair operator sequence with a characteristic inverted repeat sequence (29, 30). Using the FapR operator from the *fabHA-fabF* regulatory region, we used electrophoretic mobility shift assays (EMSA) to compare the response of purified FapR and FapR* to malonyl-CoA. When operator DNA was incubated with FapR, nearly all operator-containing fragments were shifted, whereas the control DNA fragment was unchanged. Upon titration with increasing concentrations of malonyl-CoA, the unshifted operator DNA band re-appeared (Figure 4A), suggesting that FapR is released from the DNA in response to its ligand. In comparison, DNA-binding by FapR* was relatively insensitive to malonyl-CoA (Figure 4B). With FapR, 360 µM malonyl-CoA was sufficient for release of 50% of operator DNA, whereas even >1 mM malonyl-CoA caused only a small decrease in the amount of FapR*-DNA complex (Figure 4C). We note that FapR* protein cannot constitute a malonyl-CoA blind protein since the complete elimination of malonyl-CoA binding is lethal (30). Therefore, a small decrease in the percentage of bound DNA in response to high concentrations of malonyl-CoA is consistent with a protein that still responds to malonyl-CoA, albeit with substantially decreased affinity. These results support our hypothesis that FapR* is a super-repressor that decreases the expression of FAS enzymes.

**Figure 4.**
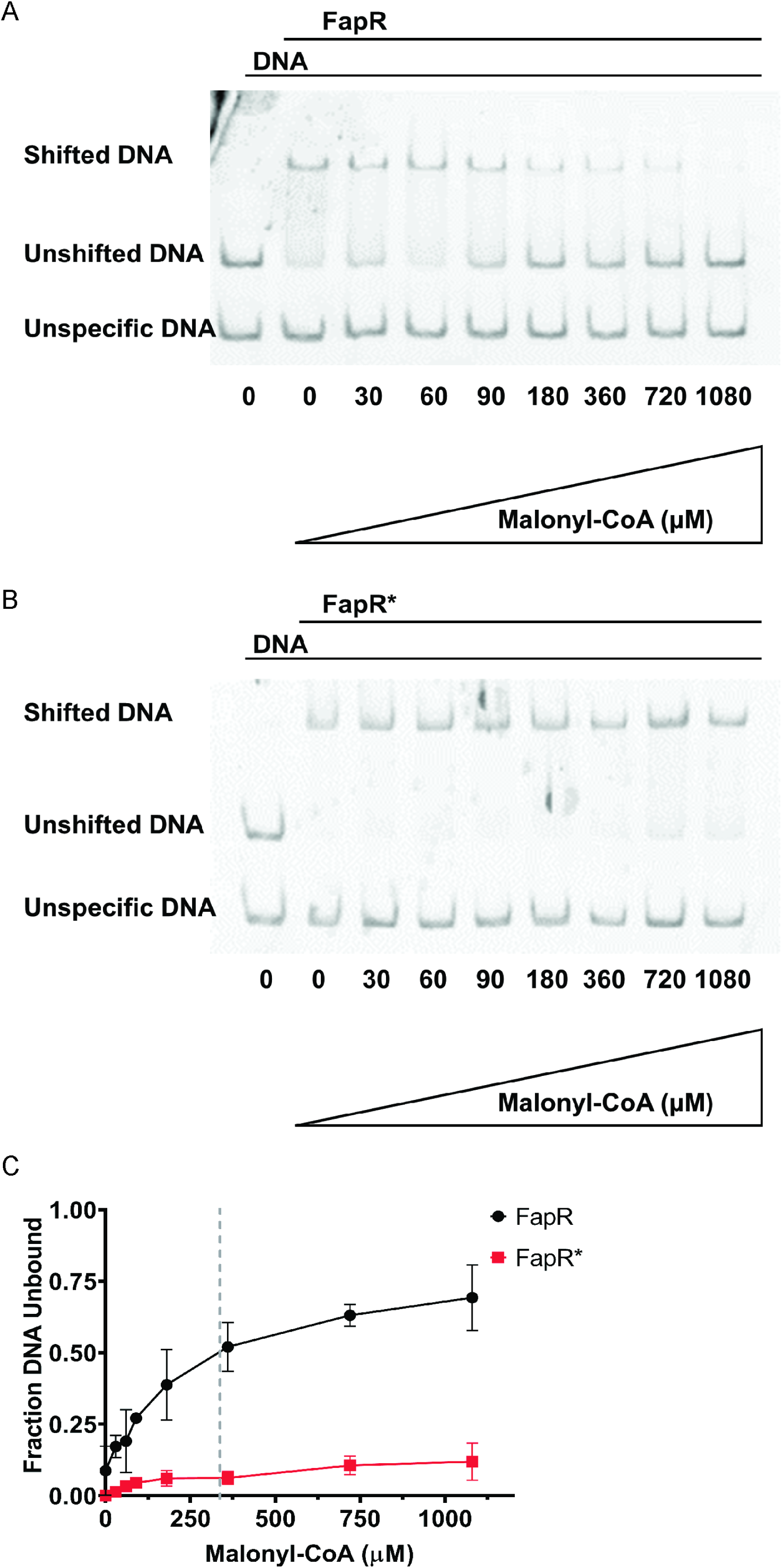
FapR* exhibits altered interactions with malonyl-CoA. (A) Electrophoretic mobility shift assay (EMSA) where a FapR-DNA complex was titrated with increasing concentrations of malonyl-CoA. The lower DNA bands serve both as a loading control and control for protein interaction specificity as it does not contain the FapR-recognized inverted repeat and is therefore not expected to shift upon addition of FapR. The ratio of protein-DNA was kept constant for all conditions. (B) EMSA with FapR*-DNA complexes titrated with malonyl-CoA under the same conditions as above. (C) The fraction unbound DNA with different concentrations of malonyl-CoA was determined by calculating the ratio of intensity of the shifted protein-DNA band to the unshifted control band. All values were normalized to 1. The dotted line represents the concentration of malonyl-CoA where 50% of the DNA is unbound by FapR. N = 3 trials.

### FapR* does not rescue PG-limited cells by increasing membrane fluidity

We hypothesized that altering the relative expression of genes in the FAS pathway might increase membrane fluidity, which has recently been recognized as a physical property correlated with movement of the elongasome (9, 35). To determine whether increased membrane fluidity might be advantageous for the PG-limited *sigI ponA* double mutant, we tested the effect of the chemical fluidizing agent benzyl alcohol (BnOH). Indeed, BnOH rescued growth of *sigI ponA* at 37 °C on LB medium in the absence of Mg^2+^ supplementation. We note that under these conditions, the *sigI ponA* strain had a high propensity for generating suppressor mutations (Figure 5A). Whole genome sequencing revealed mutations in *walH,* a negative effector of the essential WalK-WalR two-component system (Table S2), consistent with our previous finding that gain-of-function mutations in WalK rescues PG-limited cells (19). In contrast, the *rasP ponA* strain could not be rescued by BnOH under the same growth conditions (Figure 5A), suggesting that the RasP-dependent σ^W^ stress response, which is induced by membrane fluidizing agents (36), may be important for rescue. Consistently, a *sigI ponA sigW* triple mutant was less responsive to BnOH than a *sigI ponA* strain (Figure 5B).

**Figure 5.**
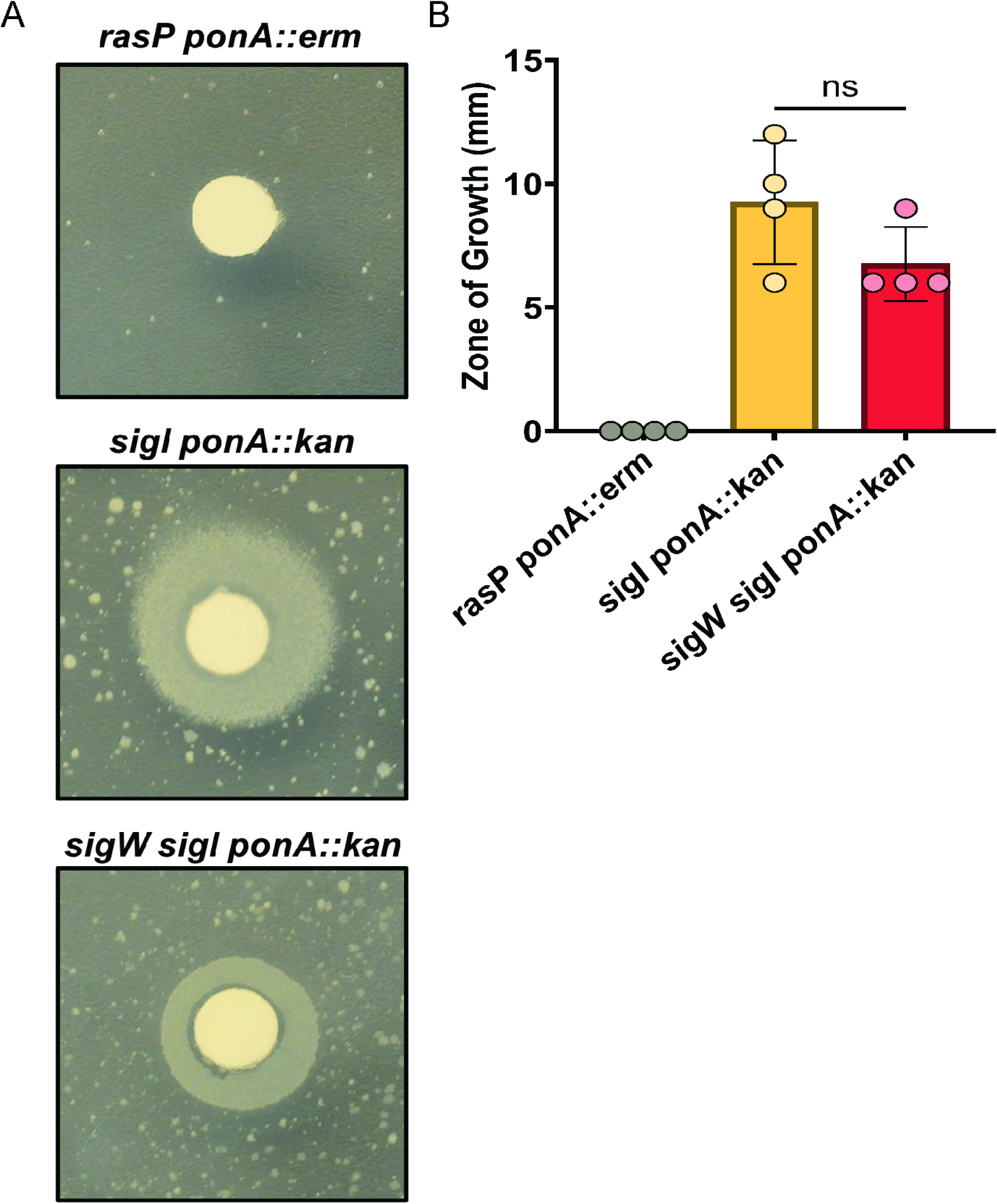
Chemical rescue of PG-limited cells by benzyl alcohol (BnOH). (A) A representative zone of growth obtained following application of 10% BnOH to an LB plate without Mg^2+^, inoculated with the indicated strains. *sigI ponA* and *sigW sigI ponA* are able to grow near the BnOH disc, with numerous suppressor mutations also apparent. The *rasP ponA* strain is unable to be rescued by BnOH, and forms fewer suppressor mutations. (B) Quantitation of the zone of growth on LB medium. A one-way ANOVA with Tukey test for multiple comparisons was performed. *p*-value > 0.05. The size of the filter disc (6 mm) was subtracted from the zone of growth values.

To directly assay for changes in membrane fluidity in the *fapR\** mutant, we employed fluorescence anisotropy using the probe 1,6-diphenyl-1,3,5-hexatriene (DPH). We observed a small, but statistically significant increase in anisotropy (a lower degree of rotational freedom) (Table 2). In contrast to our expectation, this implies that the *fapR*\* mutant has a small decrease in membrane fluidity, rather than an increase as typically seen with BnOH (37). The observed decrease in membrane fluidity in the *fapR*\* strain is supported by measurements of the fatty acyl composition of the membrane (Table 2). FapR* cells contained a significant increase in the ratio of C_17_ to C_15_ acyl-chains, with longer acyl chains known to be correlated with decreased fluidity (38). We conclude that an increase in fluidity can benefit PG-limited cells, as supported by the BnOH results, but this is unlikely to be related to the rescue of growth seen in the presence of *fapR\**.

**Table 2.**
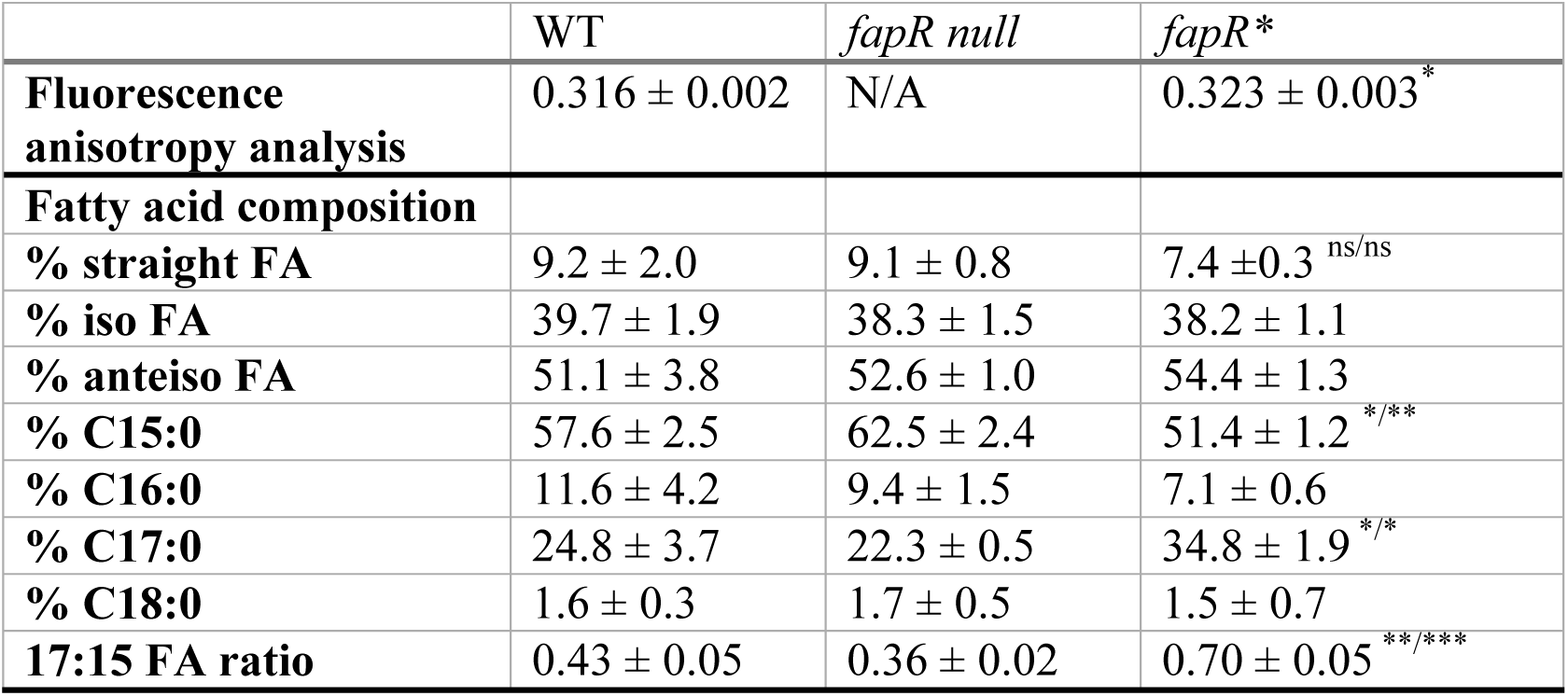
The fapR* strain has an altered membrane composition. Data derived from fluorescence anisotropy analysis of WT and fapR*. The data presented represents the average of three biological replicates where “±” represents the standard error. A Student’s t-test was performed and the anisotropy of WT and fapR* were found to be statistically different from each other (p-value < 0.05). Data derived from FAME analysis of the following strains: WT, fapR null and fapR* membranes. The data presented represents the average of three biological replicates where “±” represents the standard error. A one-way ANOVA with a Tukey test for multiple comparisons was performed. p-value cutoffs (*) < 0.05, (**) < 0.002, (***) < 0.0002 represented as value^x/y^ where “x” indicates p-value compared to WT and “y” indicates p- value compared to fapR null.

### FapR* does not rescue PG-limited cells by increasing PG synthesis

Although *fapR*\* did not restore PG synthesis by increasing membrane fluidity (Table 2), it remains possible that it could alter membrane properties in other ways that help increase elongasome function. Therefore, we quantified the dry weight of PG from extracted sacculi (Figure 6). As the *rasP ponA* mutant was unable to grow without Mg^2+^ supplementation, we chose to grow all strains with Mg^2+^ supplementation for consistency. Even with added Mg^2+^, the *rasP* and *ponA* single mutants, and the conditionally lethal double mutant *rasP ponA*, exhibited significantly reduced levels of PG compared to WT (Figure 6). This reduced level of PG sacculus per cell mass is consistent with the inability of Mg^2+^ to completely suppress the morphological defects of the *rasP ponA* strain, as noted above (Figure S3B). Although the *rasP ponA* mutant had similar levels of PG compared to the individual *rasP* and *ponA* mutants, the time required to reach an OD_600_ of 0.4 was significantly longer. Strains containing only the *fapR*\* mutation also had significantly reduced levels of PG compared to WT cells, suggesting that this mutation reduces PG synthesis even in cells containing a full complement of aPBPs, the elongasome machinery, and intact CESR pathways. Further, we note that *fapR*\* was unable to increase the proportion of PG (relative to cell mass) in the *rasP ponA fapR*\* strain compared to the PG-limited *rasP ponA* strain (Figure 6). Thus, we conclude that *fapR*\* does not rescue growth of PG-limited cells by increasing PG synthesis. Instead, the major effect of *fapR*\* is to reduce the rate of membrane synthesis (and therefore growth rate), and under these conditions cells can tolerate a lower level of PG per cell mass.

**Figure 6.**
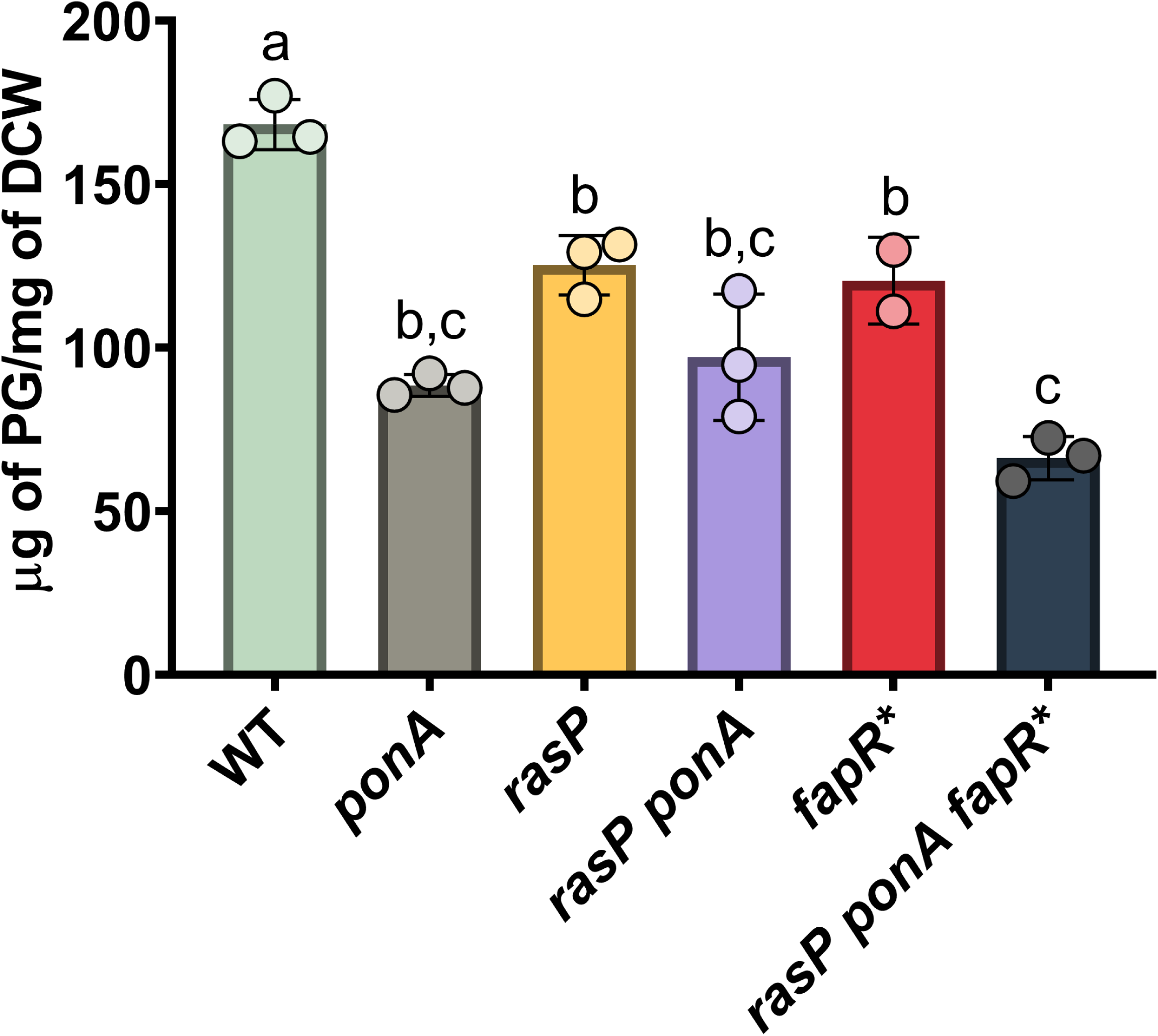
*fapR\** does not restore PG synthesis to PG-limited cells. Weight of the PG sacculus (µg of PG per mg of dry cell weight; DCW) for WT, *rasP, ponA, fapR* rasP ponA* and *rasP ponA fapR\** strains grown in LB with 10 mM Mg^2+^. A one-way ANOVA with Tukey correction was done between all strains. Columns labeled with different letters are statistically distinct from each other with *p*-value cut-off < 0.01. A “b,c” indicates that that column is not statistically distinct from any column labeled with either “b” or “c.”

### Chemical inhibition of FAS by cerulenin can rescue growth of a PG-limited strain

Since *fapR*\* rescues growth of PG-limited strains by restricting FAS, we hypothesized that antibiotics that inhibit FAS might also rescue growth. Remarkably, CER rescued growth of the PG-limited *rasP ponA* strain (Figure 7A). However, treatment with 1.65 or 3.3 µg/mL CER does not resolve the abnormal morphology of *rasP ponA* (Figure S5A), and echoing *fapR*\*, exacerbates the coiled morphology rather that restoring normal morphology (Figure S3C). With optimal concentrations of CER (1.65 or 3.3 µg/mL), growth was comparable to that of suppressed strain carrying the *fapR\** allele (Figure 7A).

**Figure 7.**
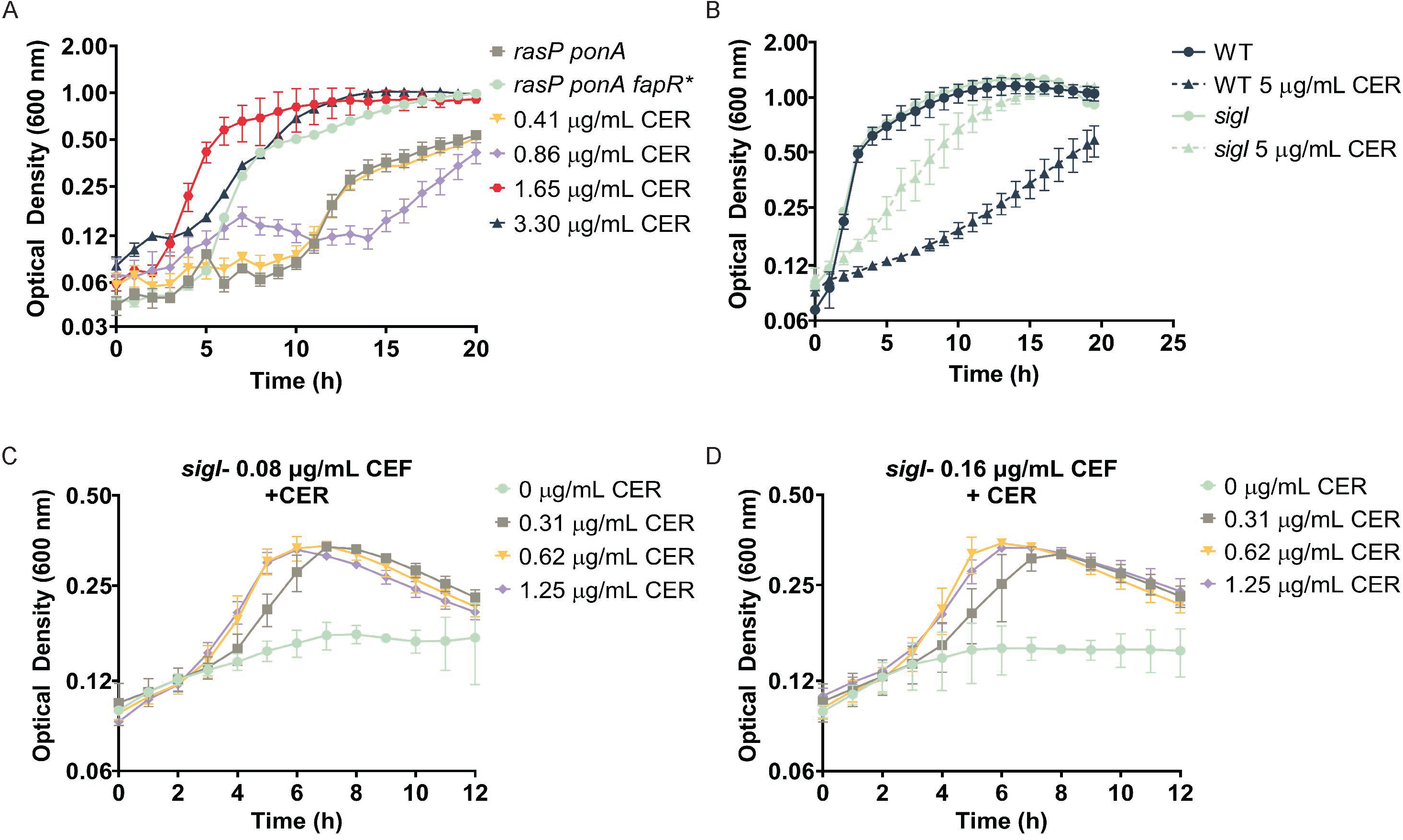
Restriction of membrane synthesis enhances growth of PG-limited cells. (A) Increasing concentrations of CER rescue growth of the *rasP ponA* strain at least as well as *fapR\** when grown in LB medium without Mg^2+^. (B) The *sigI* null mutant is limited in its ability to upregulate PG synthesis and has a higher tolerance for CER (5 µg/mL) than WT. (C-D) The *sigI* strain is known to be CEF sensitive due to an inability to up-regulate elongasome function when aPBPs are inhibited. Treatment of *sigI* with sub-MIC levels of CER increases *sigI* tolerance to CEF, demonstrating an antagonistic relationship for these two antibiotics.

Previously, CER was found to reduce protein synthesis similar to that experienced following nutrient deprivation (39, 40). However, PG-limited cells could not be rescued by treatment with sub-MIC levels of the protein synthesis inhibiter chloramphenicol (Figure S5B), or by reducing growth rate through growth on non-preferred carbon sources (Figure S5C). Synthesis of the second messenger (p)ppGpp has also been implicated in adaptation to fatty acid starvation (41). However, despite increased FAS repression in our strains, rescue by *fapR*\* is independent of (p)ppGpp. Specifically, we demonstrate that a *sigI* null mutant, which is sensitive to the β-lactam antibiotic cefuroxime (CEF), is suppressed by *fapR\** independently of (p)ppGpp (Figure S6).

Although *sigI* mutants are very sensitive to CEF (Figure S6), they have an increased resistance to CER compared to WT cells (Figure 7B). Further, growth in the presence of otherwise inhibitory concentration of CEF can be rescued by CER (Figure 7C, 7D). Thus, a reduction in membrane synthesis by either *fapR\** or CER is sufficient to restore growth of PG-limited cells (*rasP ponA*) and *sigI* cells defective in the ability to upregulate PG synthesis in response to cell envelope stress (Figure 7). Consistently, the *rasP* single mutant, despite being more sensitive to both CEF and CER than the *sigI* mutant, also displayed a notable antagonistic relationship between the two antibiotics. The *rasP* mutant cells inhibited in growth by CEF could be rescued by low levels of CER (Figure S7).

## Discussion

*B. subtilis* is an excellent model system for defining how cells synthesize a cell envelope during growth and cell division. *B. subtilis* has a relatively simple envelope comprising a cell membrane and peptidoglycan layer and can be grown in a wide variety of chemically defined media. Considerable progress has been made in defining how synthesis of each individual envelope layer is regulated. For example, in *B. subtilis,* FAS and phospholipid synthesis are coordinated in response to availability of the malonyl-CoA precursor, which binds as an inducer to the FapR repressor (30). Depletion of the fatty acyl-phosphate synthase, PlsX, halts both FAS and phospholipid synthesis, which suggests that *plsX* also plays a key regulatory role in coordinating FAS and phospholipid synthesis (42, 43). Similarly, PG synthesis is homeostatically regulated by multiple mechanisms that control the GlmS-dependent partition of fructose 6-phosphate into amino sugar synthesis. GlmS expression is feedback regulated by a GlcNAc-sensing riboswitch and by UDP-GlcNAc inhibition of GlmR, an activator of GlmS activity (44, 45). The later stages of PG synthesis are feedback regulated by lipid II availability through a pathway involving the membrane-embedded kinase PrkC, which modulates the number of active elongasome complexes in the cell through phosphorylation of RodZ and possibly other proteins (46).

Although membrane and PG synthesis generally occur at appropriate rates to sustain cell growth, disruptions in their coordination can occur in response to chemical stress, including antibiotics that impair FAS or PG synthesis (16, 47), or physical stress such as osmotic shifts (48). Perturbations of cell envelope synthesis often induce stress responses coordinated by specific cell envelope stress response (CESR) pathways (Figure 1). For example, impairment of PG synthesis leads to the induction of PG synthesis enzymes, and alternate enzymes to compensate for those that might be rendered inactive by antibiotics. However, there is no described mechanism for impaired PG synthesis to down-regulate membrane synthesis. Further, synthesis of the cell membrane and PG layer must be uncoupled under some conditions. For example, L-forms are able to synthesize membranes even in the absence of PG (49–51), and photosynthetic bacteria can synthesize abundant internal membranes without a coordinate increase in PG synthesis (52).

There is considerable evidence that altering cell membrane synthesis can have consequences for the rate of PG synthesis. In *E. coli*, FAS has been implicated as a main determinant of cell size (40). Spontaneous suppressor mutations in *fabH* can compensate for lethal defects in LPS synthesis and reduce the rate of cell envelope growth (53). Further, *B. subtilis* supplemented with exogenous fatty acids will increase in size by up to 10 percent, suggesting that increased membrane synthesis may allow increased cell wall synthesis (40). The mechanism by which membrane synthesis can alter the rate of PG synthesis is not fully understood, but may be related to the localization of PG synthesis enzymes in the cell membrane. For example, in response to hyperosmotic shock loss of water may lead to plasmolysis in which the close association of the membrane with the PG layer is locally lost. Under such a condition, PG synthesis is thought to continue but cell length no longer increases until the osmotic stress is relieved.

The question of how membrane synthesis might respond to perturbations in PG synthesis is even less well understood. Under iso-osmotic conditions, impaired PG synthesis can lead to the emergence of L-forms in which continued membrane synthesis leads to the extrusion of membrane from within the sacculus and the continued growth of cells lacking a PG wall (54). At least in *B. subtilis*, the emergence of L-forms typically requires mutations that up-regulate membrane synthesis (50, 55), together with mutations that alleviate oxidative stress (56).

We have approached the question of how cells respond to a restriction in their capacity for PG synthesis from a different perspective. As shown previously, cells lacking all four aPBPS (Δ4 aPBP) or even just the dominant aPBP (*<u>ponA</u>*) are viable on LB medium only because stress responses increase the synthetic capacity of the elongasome. The activation of σ^M^ increases expression of the RodA TG, as well as other PG synthesis enzymes, and a *ponA sigM* double mutant is inviable (6). Similarly, σ^I^ is required in a *ponA* null mutant to increase elongasome activity by increasing expression of elongasome-associated MreBH and LytE proteins (19).

Using forward genetics, we selected for suppressors of PG-limited strains lacking the major aPBP (*ponA* mutation) and defective in the σ^I^ pathway (e.g. *ecsA, rasP* or *sigI* mutants) (Figure 8). By selection for growth on LB we have identified two main mechanisms of suppression. One mechanism of rescue involves the up-regulation of the essential WalK-WalR two-component system through either gain-of-function mutations in WalK (19) or mutations affecting the negative regulator WalH (Table S2, Figure 8). The WalKR two-component system regulates the *mreBH* and *lytE* genes that would otherwise be upregulated by σ^I^.

**Figure 8.**
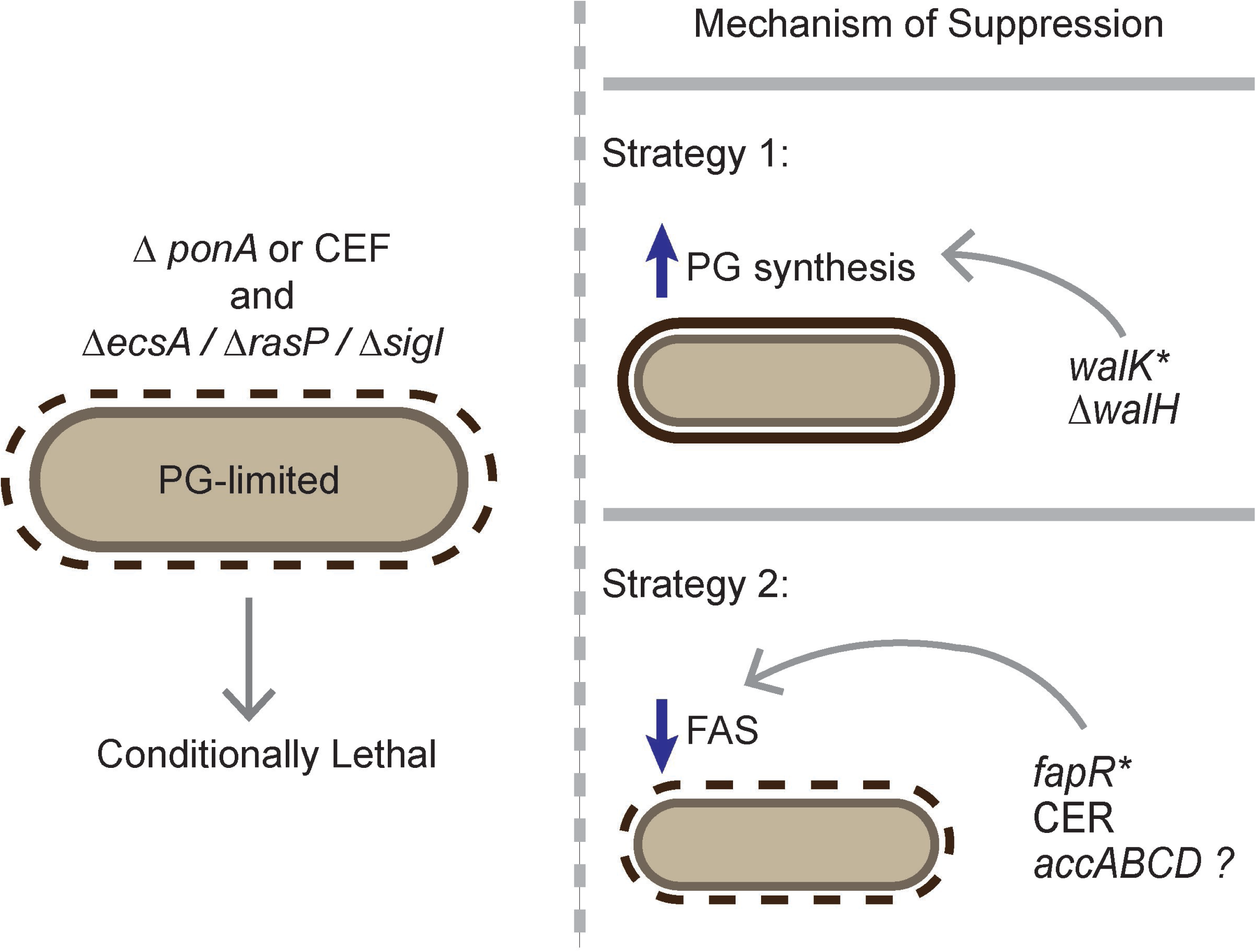
Survival strategies to restore viability of conditionally lethal PG-limited cells. Cells deficient in aPBP activity (Δ*ponA* or treatment with CEF) and with limited capacity to up- regulate the elongasome via the *ecsA, rasP, sigI* pathway are considered PG-limited and are unable to grow on LB medium in the absence of added Mg^2+^. Isolation of spontaneous suppressor mutations that restore viability to PG-limited cells reveal two mechanisms of suppression. One set of suppressor mutations affect the essential *walKR* pathway to increase elongasome-directed PG synthesis as shown previously (19), and supported by the isolation of *walH* suppressors in this study. Alternatively, we here report that a spontaneous suppressor mutation (*fapR\**) that down-regulates FAS can also rescue growth. Consistently, growth is also rescued by chemical inhibition of FAS by CER. We also recovered numerous mutations affecting the ACC complex (*accABCD*) that likely work in the same way, but we were unable to reconstruct these point mutations in a WT background for detailed characterization. Collectively these results indicate that reduced membrane synthesis is sufficient to rescue growth of PG- limited cells.

Here, we focused on a second cohort of suppressors mutations affecting FAS (Figure 8). We recovered missense mutations that we hypothesize decrease ACC activity. However, efforts to further explore these mutations were frustrated by our inability to introduce these mutations into WT cells. This hints that an insufficient capacity for FAS is not well tolerated in cells with normal PG synthesis capacity. YqhY has been implicated as a possible negative regulator of ACC (57, 58). However, induction of *yqhY* did not rescue our PG-limited strains (data not shown), suggesting that simply overexpressing the already abundant YqhY protein may not be sufficient to downregulate ACC activity.

We also recovered several mutations affecting FapR, a repressor of FAS that responds to malonyl-CoA as inducer. Detailed characterization of one suppressor, *fapR*\*, demonstrated that this mutation results in a FapR* super-repressor that decreases the rate of FAS leading to slower growth and an ability of cells to tolerate the reduction in PG. These findings suggest that PG-limited strains are unable to form colonies on LB due to stresses imposed by an excess capacity of membrane synthesis. This may lead to membrane extrusion or blebbing, similar to what occurs during L-form emergence (54), and under these conditions this may lead instead to cell lysis and death. This further implies that a limitation in PG synthesis, here imposed genetically, does not automatically trigger a reduction of membrane synthesis. Consistent with the idea that PG-limited cells can be rescued by a reduction in membrane synthesis, CER can also restore growth of the *rasP ponA* mutant (Figure 7A).

Collectively, our results suggest that a limitation in PG synthesis leads to substantial cell stress that can lead to diverse outcomes. Up-regulation of CESR pathways can help restore PG synthesis to facilitate survival (16, 47). Alternatively, cells may transition to an L-form like state that allows growth in the absence of a cell wall, although at least in *B. subtilis* this requires an iso-osmotic environment and additional facilitating mutations (50, 55). Finally, as shown here down-regulation of FAS may relieve the stress sufficiently for cells to persist despite having a reduced ratio of PG to cell mass. A more complete understanding of the balance between FAS and cell wall synthesis may provide further insights relevant to antimicrobial treatments and the maintenance of cell homeostasis.

## Materials and Methods

### Growth conditions, bacterial strains, and plasmids

All strains were cultured in lysogeny broth (LB) medium at 37 °C and aerated on an orbital shaker at 300 RPM. Glycerol stocks were streaked on LB agar plates and grown overnight at 37 °C. When needed, antibiotics were used in growth media at the following concentrations: 100 µg/mL ampicillin; 10 µg/mL chloramphenicol; MLS (1 µg/mL erythromycin plus 2.5 µg/mL lincomycin); 10 µg/mL kanamycin; 100 µg/mL spectinomycin, 10 µg/mL tetracycline. All synthetic lethal strains were grown in media supplemented with 20 mM MgSO_4_. Bacterial strains used in this study are listed in supplemental Table S3. Deletion strains were created by utilizing the BKK/BKE genomic knockout library available at the Bacillus Genome Stock Center (BGSC). Gene deletions with kanamycin or erythromycin cassettes were moved into the lab WT 168 strain, taking advantage of the natural competence of *B. subtilis* grown to OD_600nm_ 0.8 in modified competence (MC) media. Null mutations were created by removing the antibiotic resistance cassette to create a clean in-frame deletion mutation using pDR244 as previously described (Koo et al., 2017). Mutations were confirmed via colony PCR using the designated check primers as detailed in Table S3. For strains with low levels of natural competence, SPP1 transfection was used to create mutations as preciously described (59). The FapR* suppressor mutations were reconstructed via CRISPR-cas9 mutagenesis using the pJOE8999 as described previously (60). Primer sequences for repair template, guide RNA and construct confirmation can be found in Table S3. Constructs produced via CRISPR-cas9 mutagenesis were confirmed via Sanger sequencing performed by the Cornell University Biotechnology Resource Center (BRC). Genes were expressed ectopically at the *amyE* locus using the pPL82 plasmid (61) and expressed from a P_s*pac(hy)*_ promoter after induction with 1 mM IPTG.

### Plating efficiency

All strains were grown in 5 mL LB supplemented with 20 mM MgSO_4_ at 37 °C with shaking until OD_600nm_ ∼0.4-0.5. The cells were washed by centrifuging 1 mL of culture at 5000 RPM, and the pellet was resuspended in 1 mL LB without additional Mg^2+^. Serial dilutions were performed and 10 µL of culture was spotted onto an LB-agar plate, air dried for 15 minutes and then incubated overnight at 37 °C. N = 3. A representative image is shown.

### Growth assay

From a colony, strains were grown in 5 mL LB medium supplemented with 20 mM MgSO_4_ at 37 °C with shaking until OD_600nm_ ∼0.4-0.5. The cells were washed by centrifuging 1 mL of culture at 5000 RPM, and the pellet was resuspended in 1 mL LB medium to remove traces of MgSO_4_ or resuspended in minimal media (15 mM (NH_4_)2SO_4_, 0.8 mM MgSO_4_ 7H_2_O, 3.4 mM sodium citrate dihydrate, 2 mM KPO_4_, 4.2 mM potassium glutamate, 40 mM morpholinepropanesulfonic acid (MOPS), pH 7.4, 0.25 mM Tryptophan, 5 µM FeSO_4_, 5 µM MnCl_2_, 2 % carbon as specified) as required. 1 µL of this culture was added to 199 µL of media with the desired drug concentration into the well of a 100-well Honeycomb plate (Steri). The OD_600nm_ of each well was measured at 37 °C with shaking in a Bioscreen C growth curve analyzer (Growth Curves USA, NJ) every 15 minutes up to a maximum time of 24 hours. All data shown is representative plots and standard deviations from three biological replicates.

### Disc diffusion assay

Cultures were grown in LB medium at 37 °C with aeration and synthetic lethal cultures were supplemented with 20 mM MgSO_4_. Cells were harvested at ∼0.4 OD_600nm_, and where applicable, pelleted and resuspended in fresh LB medium to remove traces of MgSO_4_. 100 µL of the liquid culture was added to 4 mL of top agar (0.75 % agar) maintained at 50 °C to prevent solidification. After mixing, top agar and cells were poured onto a 15 mL agar plate (1.5 % agar) and allowed to solidify. Top agar was allowed to dry for 15 minutes prior to placing a 6 mm diameter Whatman paper filter disc on the surface of the top agar. The compound of interest was them immediately applied to the filter disc and the plates were incubated overnight at 37 °C. The diameter of the zone of inhibition or the diameter of the zone of growth were measured and recorded. Compounds were used at the following concentrations: 10 µL of a 10 % solution benzyl alcohol (BnOH), 10 µg cerulenin, 5 µg triclosan unless otherwise specified, 0.8 µg erythromycin, 8 µg chloramphenicol, 10 µg cefuroxime.

### Transcript analysis (Real-time RT-PCR)

To determine gene expression via RT-PCR, strains were grown in LB medium to ∼OD_600nm_ 0.4. To extract RNA, 1.5 mL of cells were incubated at 37 °C for 15 minutes in lysozyme-TE (30 mM Tris-HCl, 1 mM EDTA; pH 8.0 with 15 mg/mL lysozyme) to mediate cell lysis. RNA was purified as per the manufacturer’s instructions using the Qiagen RNeasy Kit. The extracted RNA was DNase treated according to the Turbo DNase kit manufacturer’s instructions (Ambion). 2 µg of RNA was used in a 20 µL reaction to synthesize 100 ng/µL of cDNA using the Applied Biosystems high-capacity reverse transcription kit. Gene expression was measured using 100 ng of cDNA and 0.5 µM of gene specific primer (see Table S3) and 1x SYBR (Bio-Rad) in a qPCR QuantStudio 7 Pro (ThermoFischer Scientific). *gyrA* was used as a house-keeping gene. Gene expression was normalized to *gyrA* and then expression values (2^-Δct^) were recorded for each gene.

### Purification of FapR and FapR*

*E. coli* BL21 DE3 cells were freshly transformed with a pET-16b expression vector containing an IPTG inducible copy of *fapR* or *fapR\** with an N-terminal 10x-His tag. Transformants were grown overnight in LB medium supplemented with ampicillin as specified and 1 % w/v glucose. The next day 100 µL of the overnight culture was used to inoculate a 3 mL pre-culture of LB medium with 1 % glucose and ampicillin. The pre-culture was used to inoculate 500 mL of LB medium with 1 % glucose and ampicillin and allowed to grow until ∼0.2 OD_600nm_. Cultures were induced with 1 mM IPTG and additional ampicillin and left at 37 °C for 3 hours. Cultures were spun down in a chilled rotor at 3500 RPM for 20 minutes at 4 °C and pellets were stored at −80 °C until purification. Cells pellets were resuspended in 5 mL chilled resuspension buffer (20 mM Tris-HCl, 500 M NaCl, 10 mM Imidazole, 0.5 tablet of cOmplete Mini protease inhibitor cocktail (Roche), pH 7.4). Cells were lysed via sonication and then spun down at 4 °C at 12,000 RPM for 30 minutes. The supernatant was further processed using a 1 mL HisTrap HP column (Cytiva), prepared according to the manufacturer’s directions, washed with wash buffer (20 mM Tris-HCl, 500 mM NaCl, 30 mM imidazole, pH 7.4) and eluted with elution buffer (20 mM Tris-HCl, 500 mM NaCl, 500 mM imidazole, pH 7.4). The concentration of the purified protein was determined by Bradford assay with a BSA standard (62).

### Electrophoretic mobility shift assay (EMSA)

Labeled DNA was prepared using FAM conjugated oligos (IDT) to amplify a portion of the *pfabHAF* region (see Table S3 for primers) and the resulting amplicon was digested with *ndeI* and purified to produce FAM fragments either with or without the *fapR* binding consensus sequence. Equimolar amounts of protein and DNA were added to 10 µL reactions prepared using buffer as detailed in Schujman et al., 2003 (29). Briefly, the buffer consisted of 20 mM Tris-HCl (pH 8.0), 0.5 mM EDTA, 50 mM NaCl, 1 mM DTT, 5 % glycerol. The reactions were incubated in the dark for 10 minutes at 37 °C before addition of malonyl-CoA followed by an additional 10-minute incubation period under the same conditions. The samples were loaded onto a pre-run 10 % non-denaturing acrylamide gel in 1x TBE buffer. Electrophoresis was performed at 100 V and the gel was imaged using a Typhoon FLA 7000 (GE Healthcare Life Sciences) and quantification was performed using Fiji (63).

### Fatty acid methyl esterase analysis (FAME)

Freshly streaked strains were grown in LB medium overnight at 37 °C with shaking. The pre-culture was then used to inoculate 100 mL of LB medium and grown at 37 °C with shaking until 0.4 OD_600nm_. 20-30 mg of cells were harvested and stored at −80 °C. Sample preparation and FAME direct analysis was performed by Microbial ID (Newark, DE). Sample preparation, gas chromatography operations, calibration standards, and peak integration, were done with Microbial Identification System software (MIDI), using standards containing a mixture of straight chain saturated and hydroxy FAMEs. Reported values represent the average and standard deviations of biological triplicates.

### Fluorescence anisotropy

Fluorescence anisotropy was performed as described with modification (64). Briefly, 5 mL of cells were grown in LB medium at 37 °C with shaking to an OD_600_ ∼1.0. Cells were harvested, centrifuged (2500 x g for 3 minutes), and the pellets were then washed twice and resuspended in phosphate buffer (100 mM pH 7.0) to OD_600_ 0.15. A volume of cells was transferred to a microfuge tube (based on the volume of the quartz cuvette to be used for measurement). 1,6- Diphenyl-1,3,5-hexatriene (DPH) was added to a final concentration of 3.2 µM. Additionally, an unlabeled control was also prepared. Tubes were placed in a water bath, protected from light, and incubated at 30 °C for 30 minutes. Fluorescence anisotropy was performed with a PerkinElmer LS55 luminescence spectrometer (λ_ex_ = 358 nm, slit width = 10 nm; λ_em_ = 428 nm, slit width = 15 nm). A correction for fluorescence intensity of unlabeled cells was performed as described (65)

### Microscopic visualization and analysis

For length and width measurement (Figure S2), the indicated strains were grown until ∼0.4 OD_600nm_ and spotted onto a slide containing a thin pad prepared from 1% agarose. The cells were allowed to air-dry before application of a cover slip. The cells were imaged at 100x magnification with immersion oil using an Olympus BX61 microscope and images were captured using a Cooke Sensicam camera. Cell lengths and widths were determined using the Fiji (63) plugin, MicrobeJ (66). A minimum of 100 cell lengths and widths were recorded for statistical analysis. A one-way ANOVA with a Tukey test for multiple comparisons was performed; *p-*value > 0.05; “a” indicates no significant difference any condition. For visualization of cell morphology (Figure S3 and S5), *rasP ponA* and *rasP ponA fapR\** strains were grown overnight at 37 °C in LB supplemented with 20 mM MgSO_4_. 1 mL of the overnight culture was washed and resuspended in LB without Mg^2+^. 20 μL of this culture was then inoculated in 200 μL of LB with the required condition and incubated at 37 °C for 4 hours. 10 μL of these cells were placed on 0.8 % agarose pads and imaged using Leica DMi8 inverted microscope. Images were magnified using NIH ImageJ.

### Isolation and quantitation of peptidoglycan (PG) sacculus

Glycerol stocks were steaked onto LB agar plates with 20 mM Mg^2+^ and incubated overnight at 37 °C. For starter cultures, colonies were inoculated into 5 mL of LB with 20 mM Mg^2+^ and grown overnight at 37 °C. The following day, 1 mL of the overnight culture was used to inoculate 100 mL of LB with Mg^2+^ and grown up to 0.4-0.5 OD_600nm_ at 37 °C. To measure the dry cell weight (DCW), 5 mL of these cultures were harvested, washed, and dried overnight at 45 °C. The remaining 95 mL of cells were centrifuged, and the pellets were stored at −80 °C for PG isolation, as described further. The cell pellets were resuspended in 1 mL of 0.1 M Tris-HCl (pH 7.5) containing 4% SDS (wt/vol) and boiled (100 °C on a sand bath) for 2 hours. The samples were then centrifuged at 20,000 g for 30 minutes, washed twice with milli Q water and resuspended in 1 mL of 10 mM Tris-HCl buffer (pH 7.5) containing 10 mM MgSO_4_. The samples were then treated with 10 μg/mL DNase I for 30 minutes at 30 °C, followed by 20 μg/mL RNase A for 30 minutes at 30 °C, and finally with 0.4 mg/mL proteinase K (plus 1 mM EDTA) for 1 hour at 50 °C. To inactivate the enzymes, the samples were then heated at 95 °C for 10 minutes followed by centrifugation at 20,000 g for 30 minutes. They were washed once with milli-Q water and resuspended in 1 mL of 1 M HCl and incubated at 37°C for 4 hours. After acid hydrolysis, the samples were again centrifuged at 20,000 g for 30 minutes and washed three times with milli-Q water, then incubated overnight at 45 °C for air drying. The dry weight of PG was measured the next day and expressed as μg of PG per mg of DCW.

## ACKNOWLEDGEMENTS

This work was supported by the National Institutes of Health (grant R35GM122461 to JDH). The funders had no role in study design, data collection and analysis, decision to publish, or preparation of the manuscript. We acknowledge Heng Zhao for guidance with the reconstruction of *fapR\** suppressor via CRISPR-cas9 mutagenesis. We thank the Angert and Dörr labs for use of their light microscopes, Ankita Sachla, and Tobias Dörr for helpful comments.

## SUPPLEMENTAL INFORMATION

**Table S1. Pathways and Genes Relevant to this Study**

**Table S2. The *sigI ponA* strain forms suppressors in the PG-synthesis regulating *walH*.** The suppressor mutations were identified via whole genome re-sequencing in *sigI ponA* cells that were able to grow in LB medium in the absence of added Mg^2+^. The genome position was determined via comparison to the *B. subtilis* reference genome NC_000964.3.

**Table S3. Strains and primers used in this study**

Figure S1. ***fapR\** is a dominant mutation that reduces growth rate.** (A) *B. subtilis fapR\** has a longer doubling time compared to WT cells grown in LB medium (WT, 24 ± 0.03 min, *fapR*,* 39 ± 0.11 min). Data are presented as the average of three trials (± standard deviation). A Student’s t-test was performed and were found to be statistically different from each other (*p*- value ≤ 0.05). (B) Induction of *fapR\** also slows growth in a WT strain, indicating that *fapR\** is a dominant mutation.

**Figure S2. *fapR\** does not grossly impact cellular morphology.** (A) *fapR\** does not impact the length-to-width ratio. A Student’s t-test was performed; “a” indicates no significant difference between any population. (B) The presence of *fapR\** does not appear to grossly impact the cellular morphology compared to WT and *fapR* null strain morphologies. Scale bar 5 µm.

**Figure S3. Limitation of cell envelope synthesis results in altered cell morphology.** (A) Morphology of PG-limited *rasP ponA* cells after transfer to LB (with no added Mg^2+^) results in an irregular morphology containing a mixture of filamentous, curved, and coiled cells. (B) WT Morphology of *rasP ponA* is not completely restored by Mg^2+^ supplementation. (C) The suppressed *rasP ponA fapR\** strain still displays an irregular, coiled morphology (with an increased frequency of tightly coiled cells) despite its ability to grow in LB in the absence of added Mg^2+^.

Figure S4. **The *fapR\** mutation does not affect sensitivity to other antibiotics.** *fapR\** does not impact the permeability of the small hydrophobic antibiotics erythromycin (Erm) and chloramphenicol (Cm) through the plasma membrane. No comparison was performed between antibiotic groups. An unpaired Student’s t-test was performed for each antibiotic group. *p*-value > 0.05. The size of the filter disc (6 mm) was subtracted from the zone of inhibition values.

Figure S5 **Reducing growth rate is not sufficient to rescue *rasP ponA* growth or morphology.** (A) Treatment of *rasP ponA* cells with CER does not restore WT morphology and instead yields a coiled morphology similar to that of *rasP ponA fapR\**. (B) Inhibition of protein synthesis by chloramphenicol is not sufficient to rescue *rasP ponA* cells in the absence of Mg^2+^. (C) Growth in minimal media (MM) supplemented with a non-preferential carbon source is not sufficient to rescue the *rasP ponA* strain viability on the absence of high concentrations of magnesium.

Figure S6 **The rescuing effects of *fapR\** is not dependent on the presence of ppGpp.** A chemical *sigI ponA* mimic was created via treatment of a *sigI* null strain with 10 µg of CEF. An unpaired Student’s t-test was performed to compare *fapR* sigI* and *fapR* sigI ppGpp^0^* only. *p-*value > 0.05. The size of the filter disc (6 mm) was subtracted from the zone of inhibition values.

Figure S7**. *rasP* exhibits increased sensitivity to CER alone and higher tolerance to CEF in the presence of CER.** (A) *rasP* is more sensitive to 5 µg/mL of CER than WT treated with the same concentration of CER. Conversely, *sigI* is more resistant to this concentration of CER compared to WT (see Figure 7B). Despite both *rasP* and *sigI* mutants being deficient in *sigI*- mediated upregulation of the elongasome, *rasP* also has additional deficiencies in the expression of them σ^W^ regulon. CER resistance in a PG-limited cell may be, in part, mediated by expression of genes within the σ^W^ regulon. (B-C) Co-treatment of *rasP* with CEF and CER. Similar to *sigI*, the *rasP* mutant exhibited enhanced tolerance to CEF treatment in the presence of CER.

