## Supplementary material for "A decrease in fatty acid synthesis rescues cells with limited peptidoglycan synthesis capacity": SI materials

### SUPPLEMENTAL FIGURES

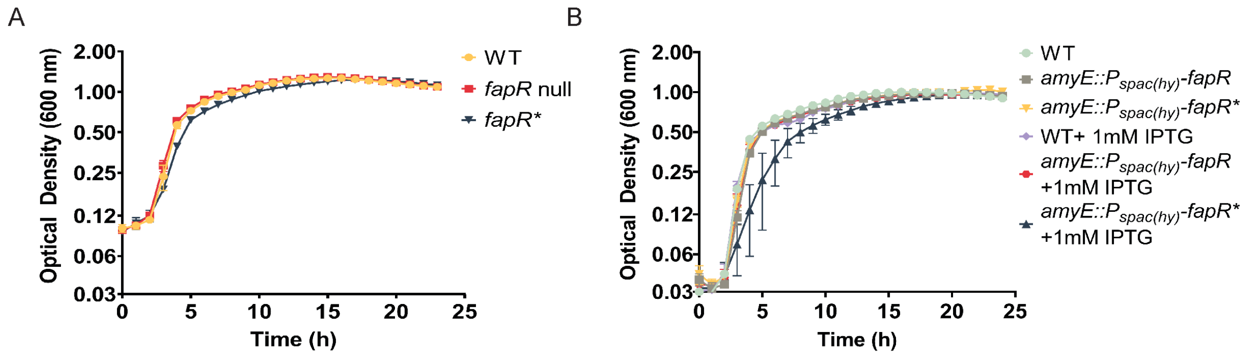

Figure S1. ***fapR* is a dominant mutation that reduces growth rate.*** (A) *B. subtilis fapR** has a longer doubling time compared to WT cells grown in LB medium (WT, 24 ± 0.03 min, *fapR*,* 39 ± 0.11 min). Data are presented as the average of three trials (± standard deviation). A Student's t-test was performed and were found to be statistically different from each other (*p*-value ≤ 0.05). (B) Induction of *fapR** also slows growth in a WT strain, indicating that *fapR** is a dominant mutation.

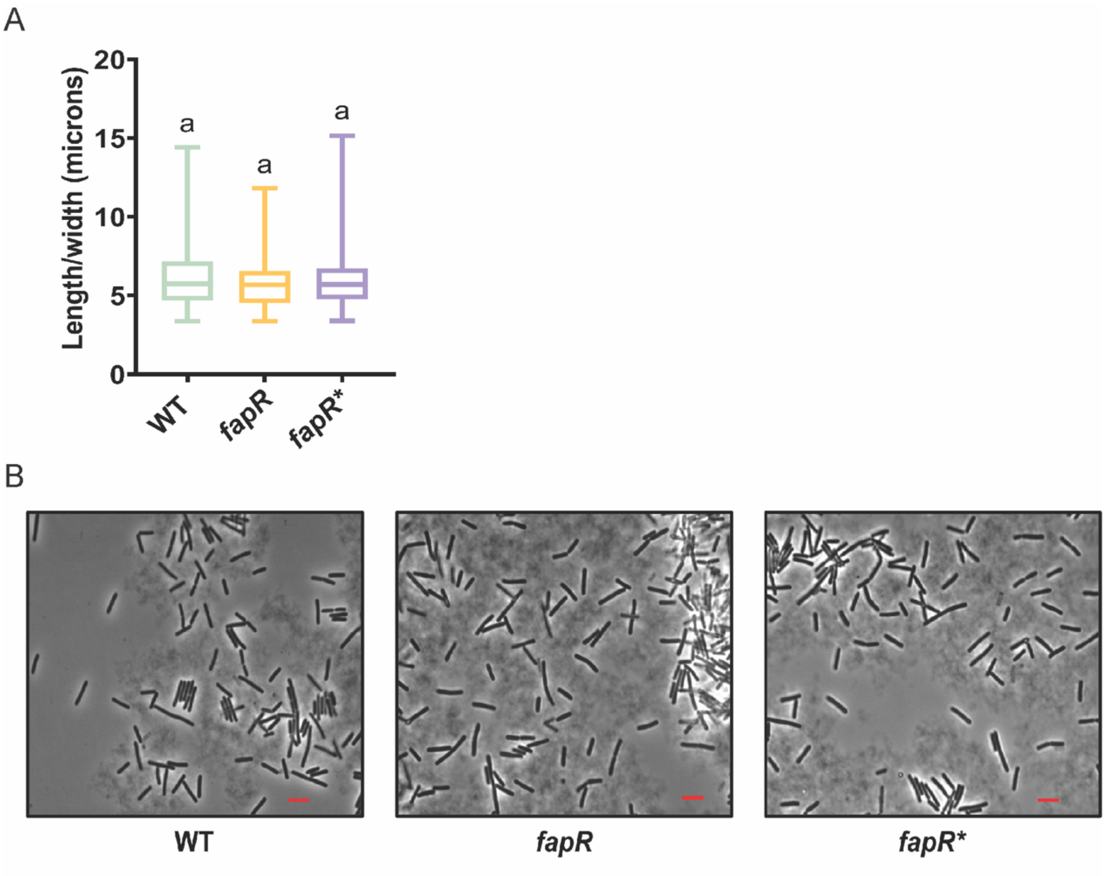

**Figure S2.** ***fapR**** **does not grossly impact cellular morphology.**

(A) *fapR** does not impact the length-to-width ratio. A Student's t-test was performed; "a" indicates no significant difference between any population. (B) The presence of *fapR** does not appear to grossly impact the cellular morphology compared to WT and *fapR* null strain morphologies. Scale bar 5 µm.

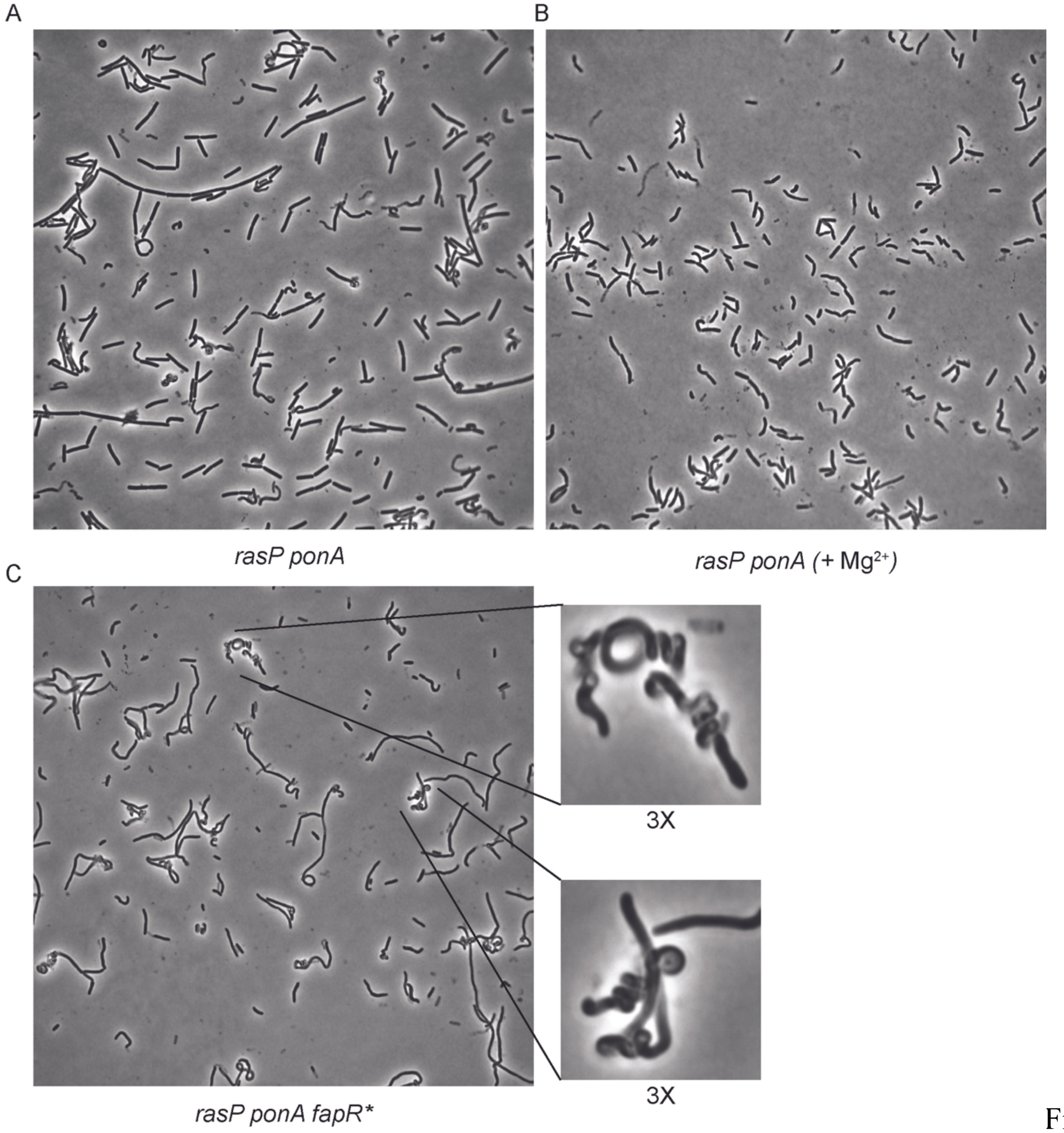

Figure S3.

**Limitation of cell envelope synthesis results in altered cell morphology.**

(A) Morphology of PG-limited *rasP ponA* cells after transfer to LB (with no added $Mg^{2+}$) results in an irregular morphology containing a mixture of filamentous, curved, and coiled cells. (B) WT Morphology of *rasP ponA* is not completely restored by $Mg^{2+}$ supplementation. (C) The suppressed *rasP ponA fapR** strain still displays an irregular, coiled morphology (with an increased frequency of tightly coiled cells) despite its ability to grow in LB in the absence of added $Mg^{2+}$.

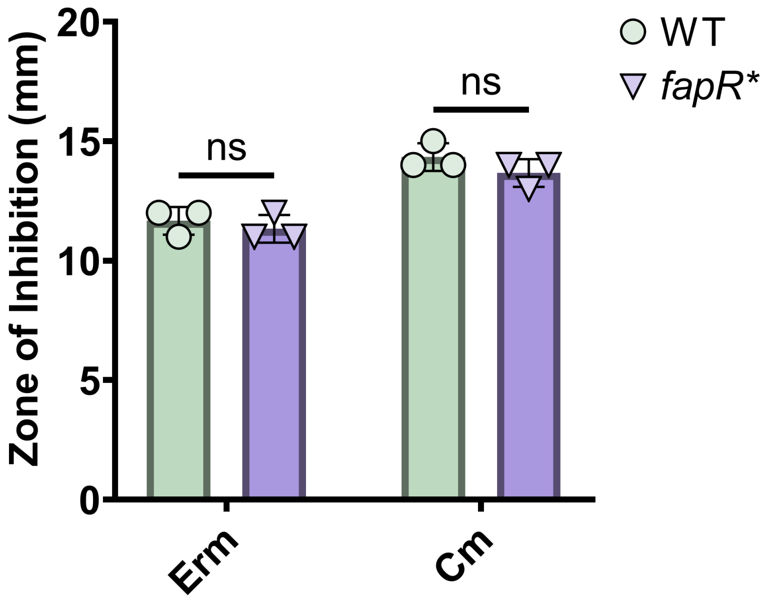

Figure S4. **The *fapR** mutation does not affect sensitivity to other antibiotics.** *fapR** does not impact the permeability of the small hydrophobic antibiotics erythromycin (Erm) and chloramphenicol (Cm) through the plasma membrane. No comparison was performed between antibiotic groups. An unpaired Student's t-test was performed for each antibiotic group. *p*-value > 0.05. The size of the filter disc (6 mm) was subtracted from the zone of inhibition values.

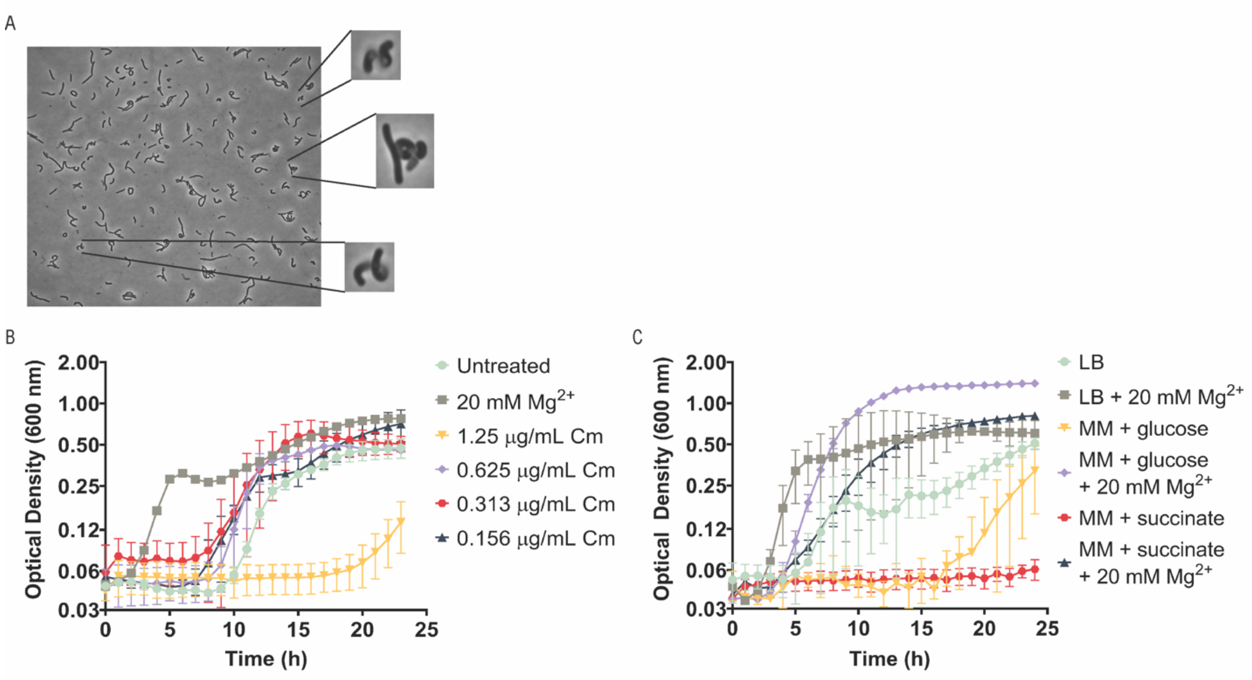

Figure S5 **Reducing growth rate is not sufficient to rescue *rasP ponA* growth or morphology.** (A) Treatment of *rasP ponA* cells with CER does not restore WT morphology and instead yields a coiled morphology similar to that of *rasP ponA fapR**. (B) Inhibition of protein synthesis by chloramphenicol is not sufficient to rescue *rasP ponA* cells in the absence of $Mg^{2+}$. (C) Growth in minimal media (MM) supplemented with a non-preferential carbon source is not sufficient to rescue the *rasP ponA* strain viability on the absence of high concentrations of magnesium.

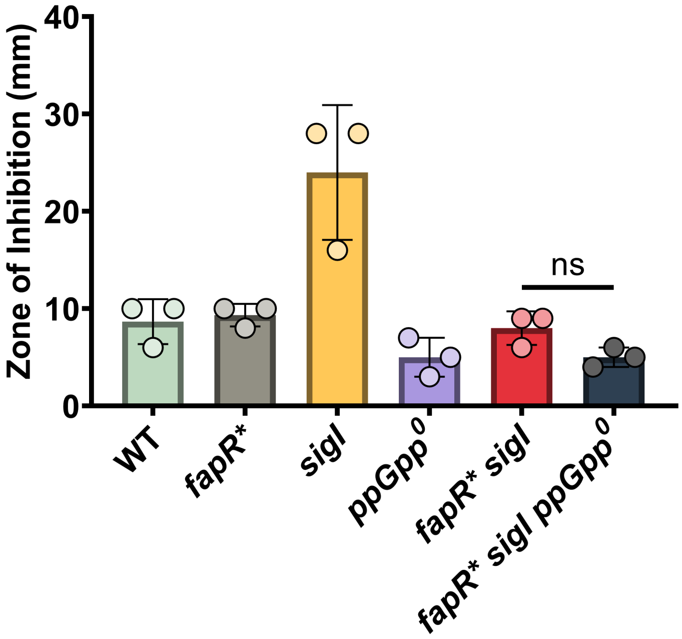

sFigure S6 **The rescuing effects of *fapR** is not dependent on the presence of ppGpp.** A chemical *sigI ponA* mimic was created via treatment of a *sigI* null strain with 10 µg of CEF. An unpaired Student's t-test was performed to compare *fapR* sigI* and *fapR* sigI ppGpp*$^0$ only. *p*-value > 0.05. The size of the filter disc (6 mm) was subtracted from the zone of inhibition values.

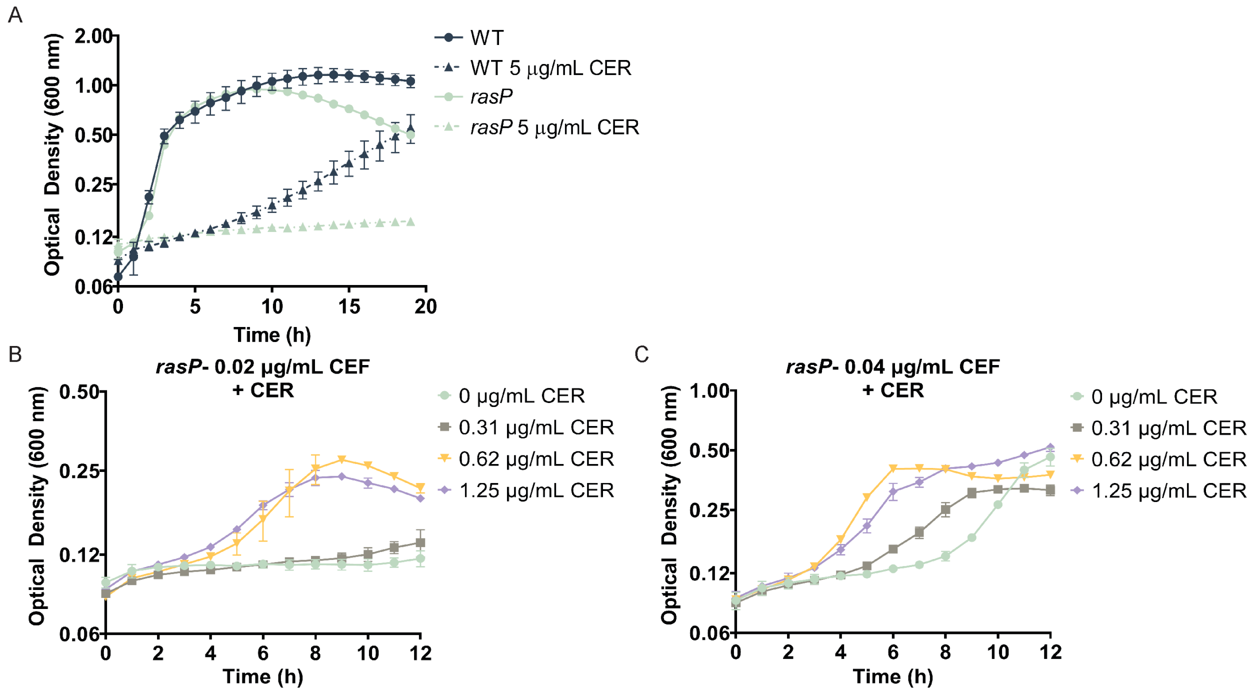

Figure S7. ***rasP* exhibits increased sensitivity to CER alone and higher tolerance to CEF in the presence of CER.** (A) *rasP* is more sensitive to 5 µg/mL of CER than WT treated with the same concentration of CER. Conversely, *sigI* is more resistant to this concentration of CER compared to WT (see Figure 7B). Despite both *rasP* and *sigI* mutants being deficient in *sigI*-mediated upregulation of the elongasome, *rasP* also has additional deficiencies in the expression of them $\sigma^W$ regulon. CER resistance in a PG-limited cell may be, in part, mediated by expression of genes within the $\sigma^W$ regulon. (B-C) Co-treatment of *rasP* with CEF and CER. Similar to *sigI*, the *rasP* mutant exhibited enhanced tolerance to CEF treatment in the presence of CER.

### SUPPLEMENTAL TABLES

### Table S1. Pathways and Genes Relevant to this Study

| Gene(s) | Function | Significance in work | Relevant References |
|---|---|---|---|
| | **Fatty Acid Synthesis (FAS)** | | |
| *fapR* | Repressor of fatty acid synthesis; de-repression elicited by binding of malonyl-CoA | Deletion of *fapR* deregulates expression of *fapR* regulon; used as a control for *fapR** phenotypes | (1) (2) |
| *fapR** | Super-repressor allele of FapR | Represses FAS to rescue PG-limited cells | This study |
| *accABCD* | acetyl-CoA carboxylase (ACC) complex; carboxylates acetyl-CoA to malonyl-CoA | suppressor mutations rescue viability of PG-limited cells | (3) and This study |
| *fabF* | FAS II condensing enzyme; elongates fatty acids by 2 carbons; negatively regulated by FapR | Target of cerulenin (CER) | (4) |
| *fabHA* | FAS II condensing enzyme; required for initiation of fatty acid synthesis in non-stressed cells; negatively regulated by FapR | Proxy for transcription of *fapR* regulon | (5) |
| | **Class A PBPs (aPBPs); PG synthesis** | | |
| *ponA* | major bifunctional glycosyltransferase/transpeptidase | Deletion is synthetic lethal in combination with either *ecsA, rasP,* or ponA | (6-8) |
| *pbpD, pbpF, and pbpG* | minor bifunctional glycosyltransferase/transpeptidase | These genes are deleted (together with *ponA*) in the Δ4 aPBP strain | (9) |
| | **Elongasome; PG synthesis** | | |
| *rodA* | SEDS family monofunctional glycosyltransferase; part of the elongasome; essential for cell wall elongation | RodA provides transglycosylase (TG) activity to the elongasome | (10, 11) |
| *pbpH, pbpH* | Class B PBPs provide transpeptidase (TP) actvity | Provide transpeptidase (TP) activity to the elongasome | (12) |
| *rodZ* | Part of the elongasome; substrate of PrkC lipid II-sensing protein kinase | May serve to regulate the density of MreB filaments and thereby growth rate | (13) |
| *mreB* | cytoskeletal actin-like protein; helps organize elongasome function during cell wall elongation | MreB functions together with other MreB paralogs (MreBH, Mbl) | (14-16) |
| *mbl* | An MreB like protein (paralog). | | (17) |

| | | | |
|---|---|---|---|
| *mreBH* | cytoskeletal actin-like protein; member of elongasome together with other MreB paralogs | Loss of *sigI* activation reduces *lytE/mreBH* expression | (6, 18) |
| *lytE* | Major cell wall autolysin; D,L-endopeptidase; regulated by SigI | Loss of *sigI* activation reduces *lytE/mreBH* expression | (6, 18, 19) |
| *mreC, mreD* | Membrane proteins functioning as part of the elongasome complex | The *mreBCD* operon is up-regulated by $\sigma^M$ | (20, 21) |
| | **WalKR: essential two-component system (TCS)** | | |
| *walK* | TCS sensor kinase; activates WalR | Mutations in *walK* rescue PG-limited cells via upregulation of *lytE* and *mreBH* | (6, 19) |
| *walR* | TCS response regulator; regulates *sigI* and autolysins including and the *mreBH/lytE* complex | WalH suppressor mutations restore viability of *sigI ponA* strains | (18) |
| *walH* | Negative regulator or WalKR system | Mutations in *walH* rescue PG-limited cells | (22) and this this study |
| | **$\sigma^M$ cell envelope stress response (CESR)** | | |
| *sigM* | ECF-type sigma factor; active in response to disruption of peptidoglycan synthesis to upregulate elongasome components and PG synthesis enzymes | $\sigma^M$ regulon includes: *ponA, mreBCD, rodA,* and lipid II flippases (*amj, yngC*), and many other genes | (20, 23) |
| | **$\sigma^I$ stress response (EcsA, RasP dependent)** | | |
| *ecsA* | ATP-binding component of an ABC transporter; essential for the function of RasP | Deletion limits elongasome activity via loss of *sigI* activation; synthetic lethal with *ponA* | (6, 24) |
| *rasP* | Site II intramembrane protease; function necessary for the cleavage of FtsL, RsgI, RsiV, RsiW | Deletion limits elongasome activity via loss of *sigI* activation; synthetic lethal with *ponA* | (6, 25, 26) |
| *rsgI* | Anti-sigma factor; cleaved by RasP and regulated by an intrinsically disordered domain | | (7) |
| *sigI* | sigma factor; control of heat shock response; regulation of elongasome | Deletion limits elongasome activity via loss of *sigI* activation; synthetic lethal with *ponA* | (6, 7) |
| | **$\sigma^W$ stress response (EcsA, RasP dependent)** | | |
| *rsiW* | Anti-sigma factor for $\sigma^W$; Cleaved by PrsW (site 1) and RasP (site 2) protease | Controls a large regulon induced primarily by membrane-associated stresses | (23, 27-29) |
| *sigW* | ECF-type sigma factor; required for adaptation to membrane active agents | Modulates membrane fluidity in response to membrane active agents via regulation of *floA/T* and *fabHAF* | (23, 30) |

**Table S2. The *sigI ponA* strain forms suppressors in the PG-synthesis regulating *walH*.** The suppressor mutations were identified via whole genome re-sequencing in *sigI ponA* cells that were able to grow in LB medium in the absence of added $Mg^{2+}$. The genome position was determined via comparison to the *B. subtilis* reference genome NC_000964.3.

| Strain Background | Gene | Genome Position | Nucleotide Change | Mutation |
|---|---|---|---|---|
| *sigI ponA* | *walH* | 4150577 | C → T | Trp429* |
| *sigI ponA* | *walH* | 4150806 | A → deletion | Val353fs |

**Table S3. Strains and primers used in this study**

| **Strains** | | | |
|---|---|---|---|
| **Strain Number** | **Genotype** | **Construction** | **Reference** |
| ***B. subtilis*** | | | |
| 168 | *trpC2* | Lab Strain | Lab stock |
| HB27097 | *trpC2 ΔecsA null ponA::erm* | Lab Strain | Patel et al., 2020 |
| HB27359 | *trpC2 ΔrasP null ponA::erm* | Lab Strain | Patel et al., 2020 |
| HB2044 | *trpC2 sigI::erm* | BGSC | Lab stock |
| HB27137 | *trpC2 ΔsigI null* | HB20406 --> pDR244 | This study |

| | | | |
|---|---|---|---|
| HB27108 | *trpC ponA::kan* | BGSC | Lab stock |
| HB27157 | *trpC2 ΔsigI null ponA::kan* | gDNA HB27157 -->HB27137 | This study |
| HB27065 | *trpC2 fapR** | CRISPR (see methods) | This study |
| HB25401 | *trpC2 ΔecsA null ponA::erm fapR** | SPP1 transduction *fapR** CRISPR plasmid -->HB27097 | This study |
| HB25402 | *trpC2 ΔrasP null ponA::erm fapR** | SPP1 transduction fapR* CRISPR plasmid --> HB27359 | This study |
| HB27139 | *trpC2 fapR* ΔsigI null* | HB27065 --> gDNA HB2044 --> pDR244 | This study |
| HB27158 | *trpC2 fapR* ΔsigI null ponA::kan* | HB27139 --> gDNA HB27108 | This study |
| HB27242 | *trpC2 fapR::erm* | BGSC | Lab stock |
| HB27064 | *trpC2 ΔfapR null* | HB27242 --> pDR244 | This study |
| HB27169 | *trpC2 ΔsigI null fapR::erm* | HB27137 --> gDNA HB27242 | This study |
| HB27207 | *trpC2 ΔsigI null fapR::erm ponA::kan* | HB27169 --> HB27108 | This study |
| HB27297 | *trpC2 amyE::*$P_{spac(hy)}$*-fapR* | 168 --> pPL82-*fapR* | This study |
| HB27298 | *trpC2 amyE::*$P_{spac(hy)}$*-fapR** | 168-->pPL82-*fapR** | This study |
| HB27214 | *trpC2 ΔsigI null amyE::*$P_{spac(hy)}$*-yqhY* | HB27137 --> pPL82-*yqhY* | This study |
| HB21116 | *trpC2 ΔsigW::erm* | Lab Strain | Lab stock |
| HB27166 | *trpC2 ΔsigW* | HB21116--> pDR244 | This study |
| HB27134 | *trpC2 ΔsigW fapR** | HB27065 --> gDNA HB21116 --> pDR244 | This study |
| HB27337 | *trpC2 ΔsigI null ΔsigW null* | HB27137 --> gDNA HB21116 --> pDR244 | This study |
| HB27338 | *trpC2 ΔsigI null ΔsigW null ponA::kan* | HB27337 --> gDNA HB27108 | This study |
| HB27173 | *trpC2 fabI::erm* | BGSC | Lab stock |
| HB27174 | *trpC2 fabL::erm* | BGSC | Lab stock |
| HB27175 | *trpC2 fabI::erm fapR** | HB27065 --> gDNA HB27173 | This study |
| HB27177 | *trpC2 fabL::erm fapR** | HB27065 --> gDNA HB27174 | This study |
| HB27230 | *trpC2 relA::erm* | BGSC | This study |
| HB27231 | *trpC2 sasA::kan sasB::tet* | gift of Heather Faega | Lab stock |
| HB27250 | *trpC2 relA::erm sasA::kan sasB::tet* (*ppGpp*$^0$) | HB27230 --> gDNA HB27231 | This study |

| | | | |
|---|---|---|---|
| HB27233 | *trpC2 ΔsigI null fapR* relA::erm* | HB27139 --> gDNA HB27230 | This study |
| HB27240 | *trpC2 ΔsigI null fapR* relA::erm sasA::kan sasB::tet (ppGpp$^{0}$)* | HB27233 --> gDNA HB27231 | This study |
| ***E. coli*** | | | |
| N/A | *BL21 DE3 pET-16b-FapR* | See Methods | This study |
| N/A | *BL21 DE3 pET16b-FapR** | See Methods | This study |
| **Primers** | | | |
| | **Primer Name** | **Sequence** | **Purpose** |
| | BKK-Kan-Check-R | GCAGGAGACATTCCTTCCGT | Cassette placement check |
| | BKE MLS check R | TTTTCTCGTTCATAGTAGTTCCTCC | Cassette placement check |
| | BKE MLS check F | CCTTAAAACATGCAGGAATTGACG | Cassette placement check |
| | ponA F | CAAGACCTCTTTCCCCCTGC | Deletion confirmation |
| | ecsA-check-F | TGATGTCCAGAACCCTGTCTC | Deletion confirmation |
| | rasP-check-F | AGTAGCTGTCGCTGCCTTTT | Deletion confirmation |
| | F-PsigI-EcoRI | ATCGAATTCCGGTTTGCGGCTGGTTTATG | Deletion confirmation |
| | SigI-Check-Rev-JRW | CTTCATAATCGCTGTTTAACGC | |
| | fapR-d8-gRNA-F | tacgAACAGGAGCACGTGTTCAGC | CRISPR guide RNA cloning |
| | fapR-d8-gRNA-R | aaacGCTGAACACGTGCTCCTGTT | |
| | fapR-d8-repair-up-F | AAGGCCAACGAGGCCGTCGGAGACCAATGACGGTT | CRISPR repair template cloning |
| | fapR-d8-repair-down-R | AAGGCCTTATTGGCCTCCCGAAACAGTCGGAAGTG | |

| | | | |
|---|---|---|---|
| | pJOE8999-check-F | CCTTTTTGCGTGTGATGCGA | CRISPR plasmid check |
| | pJOE8999-check-R | GTCAGCTAGGAGGTGACTGA | |
| | fapR-int-F | AGAAATAAGAGAGAACGCCAGGA | *fapR** sequencing confirmation |
| | fapR-int-R | TTCGCCAACGTAGCTGTTCA | |
| | fapR-check-F | GACGGTTTCGAGCTGTCTGA | Deletion confirmation |
| | fapR-check-R | ACCTCCTGCGCCATAAGAAC | |
| | fapR-HindIII-F | atcgaagcttAGTCTTAATTGTCCGGATGGT | *fapR/fapR** cloning into pPL82 |
| | fapR-XbaI-R | atcgtctagaTGCATCTACAGCTATTCTCATGCA | |
| | pPL82-check-F-HZ | AAGAAAGATATCCTAACAGCACA | pPL82 cloning check |
| | pPL82-check-R-HZ | ACGATCTTTCAGCCGACTCA | |
| | FabF-RT-Fwd-JRW | AGGAGCAAAAGGGGTGAACT | RT-PCR |
| | FabF-RT-Rev-JRW | GTGACCATCACGTCTGCATC | |
| | FapR-RT-Fwd-JRW | ATTTGCACAGGCGAATTCTT | RT-PCR |
| | FapR-RT-Rev-JRW | TTTGCTACGACACGTTCACC | |
| | fabHA-RT-F | GCTGGAATACTTGGTGTTGGAC | RT-PCR |
| | fabHA-RT-R | GCTGCAACAGCCATATGTGA | |
| | gyrA-RT-F | GGCGGCCATGCGTTATACAG | RT-PCR |
| | gyrA-RT-R | GCCATACCTACCGCAATGCC | |
| | fabI-check-F-JRW | CGCTTTTCATCTGGACAAGG | Deletion confirmation |
| | fabI-check-R-JRW | CGGGATTGCTTGATATAGATCC | |
| | fabL-check-F-JRW | CCGACATACATAACAGAAGACC | Deletion confirmation |
| | fabL-check-R-JRW | GTGCAGGTGATCGTATAGC | |
| | FapR-NdeI-pET-JRW | ATGCCATATGagaagaaataagagagaacgcc | *fapR/fapR** cloning into pET-16b |
| | fapR-BamHI-pET-JRW | ATGCGGATCCTTAttatgaatgttttgaacgatacatgtc | |
| | pET16b-Check-F-JRW | CCTCCTTTCAGCAAAAAACC | pet-16b cloning check |
| | pET16b-Check-R-JRW | GCGGATAACAATTCCCCTC | |

| | | | |
|---|---|---|---|
| | pFabHAF-L-FAM-Fwd-JRW | /56-FAM/CAAGCTCCTTTAAAGCGGG | FAM-Labeled EMSA Probe |
| | pFabHAF-L-FAM-Rev-JRW | /56-FAM/CAGATACAAGTGCTGACCG | |
| | PsigW-XbaI-F | ATCGTCTAGAACTTTGACTCCGTCATGCGT | Deletion confirmation |
| | sigW-check-R | AAGATTCGGCTGCTTGGACA | |
| | yqhY-HindIII-Fwd | atgcaagcttCGGAGGTGAATTGAATGAAAG | *yqhY* cloning into pPL82 |
| | yqhY-xbaI-Rev | atgctctagaGGTTTCGTGTTAAGCCATTTAC | |
| | relA-Check-Fwd-JRW | CGAGCTTTCTTACCTTGACG | Deletion confirmation |
| | relA-Check-Rev-JRW | GTATCATCGTGAGTGATGCC | |
| | sasA-UpCheck-JRW | CGCAAAAAGAAAGCATGGG | Deletion confirmation |
| | sasA-DownCheck-JRW | GGGCTATCAAAAGGACTTTACC | |
| | sasB-UpCheck-JRW | GAATTGCTGAAGCAGCTTTTC | Deletion confirmation |
| | sasB-DownCheck-JRW | CTTCACGATAAGAAGATCTCCC | |
